# Phenological plasticity provides limited resilience to climate warming in a sea turtle species with temperature-dependent sex determination

**DOI:** 10.64898/2026.08.06.743274

**Authors:** Blair P. Bentley, Lisa M. Komoroske, Cintia M. Santos, Armando J. B. Santos, Estefany Argueta, Vic Quennessen, Christina M. Coppenrath, Camille Kynoch, Vincent S. Saba, Claudio Bellini, Rafaely N. M. S. Ventura, J. Wilson White, Mariana M. P. B. Fuentes

## Abstract

Anthropogenic climate change is threatening global biodiversity, with sea turtles particularly vulnerable as offspring sex and developmental success are strongly influenced by incubation temperature. Behavioral plasticity, including the seasonal distribution of reproductive output, may provide short-term mechanisms for mitigating these impacts. Here, we investigated season-wide hatchling sex ratios and emergence success in a small population of green turtles (Chelonia mydas), tracking individual females across their nesting seasons. Sex ratios varied markedly through the nesting season, with later nests producing a greater proportion of male hatchlings. Moreover, sex ratios were relatively consistent among nests laid by individual females. Overall, females producing more nests over a season also produced more male offspring, suggesting that both nesting phenology and reproductive output influence individual contributions to future population demographics. Mechanistic models indicate that hatchling sex ratios have trended towards female-biased ratios (>80% female) over the past 50 years, and are projected to approach complete feminization by 2100 under continued warming. Although emergence success currently remains high (>85%), it is predicted to decline sharply after mid-century, with viable hatchling production falling to ∼30% by the end of the century. Models further show that maintaining contemporary sex ratios and emergence success will require unrealistically large delays in nesting phenology, and that even extreme shifts in phenology become ineffective by 2100. Together, these findings demonstrate that individual females can increase male hatchling production by nesting later and producing more nests, but behavioral plasticity alone is unlikely to offset the accelerating impacts of climate change on this population.

## Introduction

Climate change is expected to have widespread, adverse impacts on global biodiversity and ecosystems over the coming century, with many systems already experiencing dramatic changes (Parmesan 2006, Bellard et al. 2012, Pecl et al. 2017, Weiskopf et al. 2020). In addition to general climatic warming, climate change is altering the duration, magnitude, and frequency of extreme events such as droughts, fires, and heat waves (Reidmiller et al. 2018). Organisms such as sea turtles that possess a temperature-dependent mechanism of sex determination (i.e., the sex of the offspring is determined by the temperature experienced during incubation) are particularly vulnerable to rising temperatures and acute heat waves associated with climate change (Patrício et al. 2021, Lettrich et al. 2025). Even small changes in the nest thermal environment experienced by developing sea turtle embryos may have dramatic impacts on reproductive output, including the successful development to term, and sexual differentiation, with flow-on consequences for population demographics (Ewert et al. 1994, Mitchell and Janzen 2010, Quennessen 2026). As such, projected changes in global temperatures pose critical concerns for the persistence of sea turtle populations over the coming century (Fuentes et al. 2011), with warming of nest temperatures expected to reduce hatchling emergence success and produce increasingly female-skewed primary sex ratios (i.e., the sex ratio at the time of hatching; Hawkes et al. 2007, Patrício et al. 2021). Increasing incubation temperatures have already been observed biasing sex ratios toward females and reducing embryonic survival, with these effects documented globally across species and populations of sea turtles (Howard et al. 2014, Laloë et al. 2017, 2024, Esteban et al. 2018).

The successful development and resulting sex of sea turtle embryos are influenced by maternal choice at the time of oviposition, as the thermal environment of the nest varies across space and time (Wood and Bjorndal 2000). For example, nests laid further from the high-water mark and those lacking shading from dune vegetation generally experience relatively higher incubation temperatures than nests laid closer to the water or beneath vegetation cover. Nests laid in warmer microhabitats may therefore be more susceptible to rising temperatures (Fuentes et al. 2010b), whereas nests closer to the high-water mark may face increased risks of inundation during increasingly frequent and severe storms and tidal events (Ware et al. 2021). While spatial heterogeneity in nest conditions is relatively well documented (Wood and Bjorndal 2000; Cuevas et al. 2010; Turkozan et al. 2011; Fuentes et al. 2010a), temporal variation driven by nesting phenology is just as important, yet less well understood. Because ambient and sand temperatures are tightly correlated (Fuentes and Porter 2013; Bentley et al. 2020a), nest thermal environments vary substantially over the duration of the nesting season (Matsuzawa et al. 2002, Stokes et al. 2024). Consequently, the timing of nesting onset strongly influences the thermal conditions experienced by developing embryos oviposited in these nests.

Individual females typically lay multiple (2-12) clutches within a single nesting season, often spanning several weeks to months, covering many thermal profiles (Miller 1996). While many studies consider the impacts of sea turtle nesting phenology at the population level (Mazaris et al. 2008, Mazaris et al. 2013, Laloë and Hays 2023, Fuentes et al. 2024), relatively few have considered variation at the individual level (but see Rickwood et al. 2025), despite parental identity playing a key role in embryological development (e.g., Kynoch et al. 2024, Tezak et al. 2020a). Additionally, inter-nesting intervals may decrease with increasing temperatures, potentially compressing nesting and enabling females to lay additional clutches within a season (Hays et al. 2002; Weishampel et al. 2004), or reducing the variation in thermal environments across nests. Consequently, climatic warming may influence not only the timing of nesting onset, but also the temporal distribution and total number of clutches produced by individual females, and consequently the sex ratio of populations.

Sea turtles may respond to the impacts of rising temperatures on embryonic development by: (1) adapting the sex ratio thermal reaction norm to produce males at higher temperatures; and (2) adjusting maternal nesting behavior to select more favorable nesting habitat (Bulmer and Bull 1982). While there is evidence for local adaptation in the sex ratio thermal reaction norm in sea turtles (Bentley et al. 2020b), the unprecedented rates of climatic change and limited heritability are likely to limit the rapid evolution of this trait to keep pace with changing environmental conditions (Mitchell et al. 2013, Janzen 1994, Quennessen 2026). Consequently, plasticity in maternal nesting behaviors, such as the timing of nesting onset, will likely be required for sea turtles to continue producing nests with thermal environments that promote successful embryonic development and avoid severe male limitation in future populations (Fuentes et al. 2024).

Despite these expectations, it remains unclear how nesting phenology and the total number of clutches laid by a female interact to influence cumulative hatchling production and seasonal sex ratios (Patrício et al. 2021). If nesting is initiated earlier and more clutches are produced, individual nesting females are likely to distribute reproductive effort across a broader portion of the seasonal thermal gradient, potentially exposing clutches to a broader range of incubation temperatures, which may moderate shifts in overall primary sex ratios. Should individual females be able to alter the timing of their nesting to accommodate changing local conditions, then plasticity may provide a short-term buffer against warming-driven shifts in incubation temperatures and associated emergence success and sex ratios. (Weishampel et al. 2004). However, evidence for within-individual variation and the consequences of such within-season spreading of reproductive effort remain limited. While previous work has documented phenological shifts in response to warming climates, such temporal changes alone are unlikely to fully offset warming-driven mortality and feminization in sea turtle populations (Weishampel et al. 2004; Almpanidou et al. 2018; Monsinjon et al. 2019; Laloë and Hays 2023; Fuentes et al. 2024). Such studies have largely focused on population-level patterns of nesting phenology, with comparatively little understanding of how repeated nesting behavior interacts with phenology at the individual level (but see Rickwood et al. 2025). Moreover, individual females have been observed to produce nests with similar incubation durations (a commonly used proxy for nest thermal environment, but see Fuentes et al. 2017) across a season, suggesting that females may experience relatively similar incubation environments across the nesting season despite broader seasonal environmental variation (Reneker and Kamel 2016).

This gap in knowledge of individual-level variation persists because most studies have been unable to link multiple nests to identified females across an entire nesting season, and because nest temperatures are often measured at resolutions that fail to capture within-season and within-female variation. Currently, we lack a clear understanding of whether individual females can use phenological timing and repeated nesting to produce mixed sex ratios within a single season, thereby contributing males to a feminized population. Here, we address these limitations by extensively measuring nest temperatures in a small nesting population of green turtles (*Chelonia mydas*) at Fernando de Noronha, Brazil. We aimed to determine: (1) the degree of within-individual plasticity in emergence success and primary sex ratios across nests; (2) how contemporary emergence success and sex ratios compare with historical estimates; (3) how climate change is expected to affect emergence success and primary sex ratios; and (4) whether shifts in nesting phenology may alleviate the adverse impacts of climate change. We hypothesized that the nest thermal environment, and associated emergence success and primary sex ratios, of repeat nesters would vary across the nesting season in correlation with seasonal thermal conditions at the rookery. We further hypothesized that females producing more clutches across a nesting season would expose developing embryos to a broader range of thermal conditions, potentially buffering reductions in emergence success and increasingly female-biased sex ratios under contemporary conditions, but that this plasticity would be insufficient to offset the impacts of future climate scenarios.

## Methods

### Study site

Fernando de Noronha is a volcanic archipelago located in the Atlantic Ocean approximately 350 km north-east of the continental Brazilian coastline (3.83°S, 32.42°W). The archipelago consists of one main island and 21 smaller islands and islets, comprising a total area of 26 km^2^ (Almeida 1955, Garla et al. 2006). The majority (>70%) of the archipelago sits within a National Marine Park established in 1988, and was declared a United Nations Educational, Scientific, and Cultural Organization (UNESCO) world heritage site in 2001 in recognition of its importance for sea turtles, sea birds, cetaceans, fish, and elasmobranchs (UNESCO 2021). The islands of the Fernando de Noronha archipelago represent important feeding habitat for both green (*C. mydas*) and hawksbill (*Eretmochelys imbricata*) turtles (Bellini and Sanches 1996, Bellini et al. 2019, Bellini 2024). The main island of Fernando de Noronha also contains critical nesting habitat for a small, regional population of breeding green turtles (approximately 50 individuals per year), primarily occurring on a single beach (Praia do Leão; Bellini et al. 2026). This population has seen a steady increase in nesting numbers since their protection in 1986 (Bellini and Sanches 1996; Bellini et al. 2026), and a long-term tagging and monitoring programme has been conducted on the island by the Brazilian Sea Turtle Conservation Programme (TAMAR) since 1987 (Bellini et al. 2026). Nesting typically occurs between January and May, peaking in mid-March, and with sporadic nesting in December and June (Bellini et al. 2026). For this study, the nesting beach was patrolled nightly by 3-5 people following the first nest of the season, with empirical data collected over the duration of four nesting seasons. As the season commences with only sporadic nesting in December, and is distributed over two calendar years, we identify each season by the year with the largest density of nesting (i.e., nesting from December 2018 to May 2019 is referred to as the 2019 season). Following the convention, we collected data between the start of the 2019 season and the end of the 2023 season (hereafter Seasons 1-4 for clarity), noting that the 2020 season (Season 1) was interrupted by the global COVID-19 pandemic.

### Empirical data collection

Emerging females were encountered during regular nightly patrols, with data collected as per Bellini et al. (2026) for individual identification. Nest temperature data were collected from a total of 285 nests and 134 females at Praia do Leão over the four nesting seasons using a combination of HOBO pendant (UA-001-64; Onset Computer Corp., Bourne MA USA; accuracy: ± 0.53°C, resolution: 0.14°C), TinyTag (TGP-4017; Gemini Data Loggers, West Sussex UK; accuracy: ± 0.5°C, resolution: 0.01°C), and iButton (DS1921G; Thermochron, New South Wales AUS; accuracy ± 1.0°C; resolution: 0.5°C) temperature loggers. In all cases, temperature loggers were strategically deployed in nests to capture as much of the nesting season as possible, with the exception of Season 1 which was disrupted by the global COVID-19 pandemic (Table S1). Temperature loggers were deployed approximately halfway through oviposition to target the middle of the egg mass within the chamber. Loggers recorded at hourly intervals, and continued to log until the nest excavation process, which was conducted two days after first emergence, or after 72 total days of incubation in cases where no evidence of emergence was detected. To ensure loggers were not lost through displacement by other nesting turtles and sand movement during emergence, they were affixed to a nest marker with silk multifilament string. During nesting Seasons 3 and 4, temperature loggers were preferentially deployed in nests of females that repeatedly nested across the season, enabling assessment of thermal variation between nests of individual females across a season.

Hatchling morphometric data was collected from a subset of monitored nests (n = 323) for development rate model parameterization. To collect hatchlings for morphometrics, egg chambers were located at 45 days of incubation by following the string affixed to the nest marker, with a mesh cage (height: 80 cm with ∼45 cm buried; mesh size: 2 cm) placed around the surface of the chamber to trap emerging hatchlings. Nest cages were checked approximately every hour during regular nighttime beach patrols (∼20:00-05:00), and during the day when a rain event occurred. Emergence time was recorded when the first hatchling was sighted within the cage (typically within one hour of emergence). Following emergence, a subset of hatchlings (20-96) were randomly collected, weighed using a mass balance scale (accuracy: 0.01 g), and measured, with straight carapace length and width (SCL and SCW, respectively) and body depth (BD) recorded using vernier calipers (Dijite; accuracy: 0.02 mm). To determine if there were differences in morphometrics between seasons, generalized linear mixed models (GLMMs) were applied using the ‘lme4’ package for R (Bates et al. 2014), and tested with the anova function of ‘lmerTest’ (Kuznetsova et al. 2017). In all contrasts, GLMMs were modeled with the morphometric variable as a response variable, season as the fixed variable, and the unique nest identifier included as a random effect. Post-hoc pairwise comparisons were made between seasons using the ‘emmeans’ package (Lenth & Piaskowsk 2026). The emergence success of each nest was calculated using the methods described by Miller (1999), with individuals classified as either “alive,” “dead in nest,” or “unhatched” at the time of nest inventory. Unhatched eggs were not opened to examine the developmental stage that mortality occurred. Total emergence success was determined as the total number of alive individuals divided by the total number of egg shells and unhatched eggs within the nest chamber (Miller 1999).

We examined the relationship between nesting onset date and seasonal reproductive output using a generalized linear mixed-effects model implemented in the R package glmmTMB (Brooks et al. 2017). The total number of nests laid by each female during a nesting season was modeled as a function of nesting onset date (numeric day of year, adjusted assuming nesting commencing on the 350th day) with nesting season included as a random intercept to account for among-year variation. Because the response variable was count data and exhibited overdispersion, a negative binomial error distribution (nbinom2 parameterization) with a log-link function was used. Similarly, glmmTMB was used to model the relationship between total number of nests and the number of hatchlings produced.

### Embryological model parameterization

A population-specific embryological development rate model was generated using the EmbryoGrowth package (version 2025.12.22; Girondot and Kaska 2014) for R (R Core Team 2026). This model integrates empirical observations of nest temperature and incubation duration with hatchling morphometrics (specifically SCL) to generate temperature-dependent embryonic development rates for sea turtles. The input empirical nest temperature traces were initially trimmed such that the first data point corresponded to the timing of logger deployment within the egg chamber, and the final recording corresponded to 23:00 on the day that emergence was recorded. Where emergence data were not available, temperatures series were trimmed to 57 days from the nesting date, with these not used to build the embryological model. After removing failed recordings, half of the nest temperature traces (n=134) were used as a “training set” to parameterize the embryological model, with the remaining data (n=133) used as model validation (i.e., the “test set”). Within EmbryoGrowth, the ‘searchR’ function was used to generate a 4-parameter Schoolfield, Sharpe, and Magnuson development rate model (Schoolfield et al. 1981) using maximum likelihood. To estimate those parameters, the ‘training set’ of temperature data was used with the mean and standard deviation SCL derived from all measured hatchlings during the study period (mean SCL = 50.03 cm, S.D. = 1.97 cm), and we used the default initial parameter values (DHA=170, DHH=200, T12H=300, Rho25=385). Credibility intervals were then estimated using 10,000 Markov Chain Monte Carlo (MCMC) iterations and priors weakly informative enough to not constrain the posterior distribution (Monsinjon et al. 2022). The resulting development rate model was subsequently used to identify the putative thermosensitive period (TSP) for downstream sex ratio predictions.

While the parameters that define the thermal reaction associated with temperature-dependent sex determination (TSD) in sea turtles vary by species and population (Wibbels 2003, Bentley et al. 2020b), population-specific thresholds are currently unavailable for Brazilian green turtles. As such, we leveraged existing green turtle data provided by the ‘DatabaseTSD’ function in EmbryoGrowth. We extracted all data for green turtles where sex was available with an associated temperature from the Northern and Southern Atlantic 2023 Regional Management Unit (RMU), which resulted in input data from two previous studies in Suriname (Mrosovsky et al. 1984, Godfrey and Mrosovsky 2006). Using this data, we fit six reaction norm models (‘tsd’ function) and selected the model that best fit the data using Akaike Information Criteria for small sample sizes with the ‘compare_AICc’ function in EmbryoGrowth. Selection criteria identified the Hill and Logistic models as equally appropriate for the data, with the Logistic model subsequently used in all models of primary sex ratios.

### Predicted primary sex ratios from natural nest temperatures

The derived embryological development model was applied to empirical nest temperature data using the ‘info.nests’ function in EmbryoGrowth. The logistic TSD reaction norm was incorporated into the model to generate growth and thermal reaction norm-weighted estimates of primary sex ratio (Monsinjon et al. 2022) based on the constant temperature equivalent (CTE) generated through the temperature logger data (“TSP.GrowthWeighted.STRNWeighted.sexratio.mean”). The model was run with the “metric.end.incubation” parameter set to “observed.” Output data represented the predicted primary sex ratio (proportion male) for that nest. Sex ratios were combined with observed emergence success of the nest to predict the total number of males and females produced in that nest. Nest production was summed across females to assess total hatchling production per female per nesting season. Differences between females were assessed using a GLM within Seasons 3 and 4 (sex ratio ∼ turtle_ID), with Tukey Post hoc tests used for pairwise contrasts between individuals.

### Microclimate model of sand temperature

Previous research has demonstrated the applicability of mechanistic microclimate models for predicting sand temperatures in the context of sea turtle nesting (Fuentes and Porter 2013, Stubbs et al. 2014, Bentley et al. 2020a, Gammon et al. 2024). We used the NicheMapR microclimate model (Kearney and Porter 2017) to model sand temperatures at Fernando de Noronha outside of the period captured through empirical measurements. Our models used climate surface data from the fifth generation of ECMWF atmospheric reanalysis of the global climate (ERA5; Hersbach et al. 2020) through the ‘micro_era5’ function of NicheMapR. As described in Klinges et al. (2022), the ‘micro_era5’ function of NicheMapR leverages the mcera5 package to connect to hourly 0.27 x 0.25 degree gridded historical ERA5 data from the Copernicus Climate Change Service (C3S) at ECMWF. Hourly ERA5 climate surfaces were downloaded for every day of the year from 1978-2025. The microclimate subroutine of the function integrates the gridded surface climate data with digital elevation models (DEMs) from the elevatr package (Hollister et al. 2021) and downscales climate data with the microclima package (Maclean et al. 2019). The ERA5 climate surfaces were selected for analyses due to their high resolution, and relatively reliable estimates of air and surface temperatures (Meyer et al. 2023).

We parameterized the microclimate model using generalized physical sand properties (as described in Bentley et al. (2020a)), using the R version of the Global Aerosol Database (run.gads=2), and with the soil moisture subroutine disabled. Nest temperatures are impacted by the level of solar radiation reflected by the sand, however, no measure of sand albedo was available for the nesting beach at Fernando de Noronha. We therefore used a generalized reflectance value of 0.5 in our models.

To allow for full parameterization, the micro_era5 model was initially run for Fernando de Noronha from 1978-2025 using the ‘write_input’ parameter selected to write the input climate variables to file. These files were then used as inputs with a fully parameterized model for marine sandy beaches as per Bentley et al. (2020a) using the microclimate function of NicheMapR. This model produced hourly sand temperature estimates from 1978-01-01 00:00 to 2025-12-31 23:00 between 0 cm and 200 cm. Sand temperatures were extracted for a depth of 50 cm for downstream analyses, which is a representative depth for sea turtle nests, including green turtles (Stokes et al. 2024).

### Metabolic heating adjustments

Few predictive studies have incorporated measures of metabolic heat into nest temperature predictions (Gammon et al. 2020), despite being well documented in sea turtle nests (e.g., Broderick et al. 2001; Zbinden et al. 2006). To account for metabolic heat production on our models, we quantified the relationship between embryonic development and metabolic heating by fitting a generalized additive mixed model (GAMM) in the R package mgcv (Wood 2011). Metabolic heating, generalized as the difference between daily nest and modelled sand temperatures, was modeled as a smooth function of the proportion of total embryonic development completed (dev_prop). A cubic regression spline with shrinkage (k = 10) was used to capture the non-linear changes in metabolic heating across the duration of incubation. To account for repeated temperature measurements and nest-specific variation, nest identity was included as a random intercept effect. Models were fitted using the bam() function with fast restricted maximum likelihood (fREML) estimation.

Following GAMM model fitting, population-level predictions of metabolic heating were generated across the full range of embryonic development (0-100% development) using 1,000 evenly spaced values of developmental proportion from the model. Predictions were obtained from the fitted GAMM while excluding nest-specific random effects to estimate the average developmental heating across all nests. Standard errors and 95% confidence intervals were calculated from the model predictions. Because metabolic heating is expected to be negligible during early incubation (Gammon et al. 2020), a baseline correction was applied to predictions by subtracting the mean observed temperature difference between empirical nest temperatures and modelled sand temperatures during the first half of development from all model predictions (i.e., metabolic heating = 0 until development >50%). Predicted values below 0°C were constrained to zero to reflect that metabolic activity cannot cool a nest relative to surrounding sand temperatures. The resulting bias-corrected developmental heating curve was exported for subsequent temperature simulations.

As our GAMM modeled the relationship between proportion of development and metabolic heat, and development rate is directly linked to temperature, we developed an iterative, dynamic development model to apply metabolic heat adjustments to modelled sand temperatures. To this end, we initially used the Schoolfield, Sharpe, and Magnuson (SSM) development rate model with the parameters previously calculated in EmbryoGrowth to estimate the amount of development achieved per hour. For each nest, the total developmental requirement for hatching was estimated by integrating hourly developmental rates across the observed incubation periods from empirical nests using the SSM. This cumulative developmental value represents the number of developmental ‘units’ required to reach hatching under observed conditions. At each hourly step, the current proportion of development completed was then used to estimate metabolic heating from the fitted GAM. Predicted metabolic heating was added to the corresponding sand temperature to obtain nest temperature at that step. Developmental rate was then recalculated from the heated nest temperature using the SSM, and cumulative development was iteratively updated. The proportion of development completed was determined as the ratio of cumulative development units to the mean developmental unit requirement. Iterations were repeated until the cumulative development reached the mean developmental unit requirement, at which point incubation was considered complete and the predicted hatch date was recorded. Metabolic heat adjusted sand temperatures (i.e., synthetic nests) were calculated for every day of the year from 1978-2025 and exported for downstream analyses.

### Nest temperature model validation

To determine the robustness of our models, we used the empirical nest temperature data collected over the course of our four nesting seasons. We compared both raw microclimate sand models and metabolic heat-adjusted sand models to our empirical nest temperature observations. Initially, all data were averaged to convert hourly temperature to mean daily temperature for comparison. We then tested the fit of our models against the empirical nest data using linear regression (R^2^), root mean square error (RMSE), mean absolute error (MAE), and model bias estimates.

### Future climate projections

Unlike correlative models, which only provide reliable estimates with inputs within the range of those used to generate the model, mechanistic models use laws of thermodynamics and are able to predict temperatures outside this range (such as those expected under climate change scenarios) (Mitchell et al. 2008, Dormann et al. 2012, Kearney et al. 2014). To investigate the impacts of rising temperatures associated with anthropogenic climate change on primary sex ratios and emergence success at Fernando de Noronha, we adjusted the macroscale climate inputs using location-specific climate deltas. To do this, we averaged the macroscale climate inputs generated through the NicheMapR ‘micro_era5’ function from 2005-2025 for every day of the year. Specifically, we averaged maximum and minimum temperatures, cloud cover, wind speeds, and relative humidity, as well as daily rainfall. The resulting output was our ‘baseline year’ climate data input for adjustment. Climate deltas for ambient temperature, relative humidity, rainfall, wind speed, and solar radiation were obtained as per Saba et al. (2016), and represent a doubling of atmospheric carbon dioxide by model year 70, and then remaining fixed until model year 80 (resembling the GFDL CM2.6 models; Delworth et al. 2012). Integrating these data into the mechanistic model allowed us to make predictions of sand temperatures from 2026 to 2105 at Fernando de Noronha. Monthly climate deltas were applied to the baseline macroscale climate data, with the NicheMapR microclimate model run with the same parameters as those described above.

### Modelling sex ratios and emergence success from synthetic nests

To predict embryonic mortality and sex ratios from synthetic nest temperatures, hourly temperatures were averaged to generate daily temperatures at 50 cm depth. Daily synthetic nest temperatures were used as inputs for EmbryoGrowth using the development rate model and TSD thermal reaction norm described above for estimates from empirical nest temperatures. Estimates were generated for synthetic nests for every day of the year from January 1st 1980 to December 31st 2100. In addition, synthetic nests were produced to align with all observed nesting events over the course of the four nesting seasons, to allow for complete rookery production to be estimated. The EmbryoGrowth outputs for mean temperature during the predicted incubation period (TimeWeighted.temperature.mean) were used as input for a generalized logistic regression to estimate emergence success (as per Laloë et al. 2017). This regression assumes an upper asymptote of 86.0% emergence success, and an inflection point of 32.7° C (Laloë et al. 2017).

To generate biologically relevant estimates of total hatchling production and expected sex ratio, we developed a smoothed nesting distribution based on nesting across the study period for nesting Seasons 3 and 4 at Fernando de Noronha (Fig. S1). For our nesting distribution model, we assumed 334 nests were laid per year, with the expected number of nests per day of the year calculated from the averaged nesting distribution of Seasons 3 and 4, ranging from zero to four nests per day (Fig. S2).

We used the formulated distribution model to investigate the mitigating impacts of shifting nesting phenology under future climate scenarios. Similar to Fuentes et al. (2024), we predicted emergence success and primary sex ratios assuming a nesting phenology shift of between -360 and 360 days from present for nests laid between 2030 and 2100. In each scenario, we shifted the nesting distribution either backward or forward, but retained the same overall shape and distribution of the curve. To provide biological context, we calculated the average sex ratio and emergence success from 2005-2025 to generate ‘baseline’ metrics to compare future scenarios to.

## Results

### Nesting ecology at Fernando de Noronha

During the monitoring period over the course of four seasons (including an interruption due to the COVID-19 pandemic), a total of 1,611 interactions with nesting green turtles were recorded, with 824 of these recorded as successful nesting events. From these, 806 nests (97.8%) could be attributed to one of the 170 identified nesting turtles. Individual nesters laid an average of 4.7 nests over the course of a nesting season (mid-December to early-July; Fig. S1), ranging from between one and nine nests. Total nest number was significantly associated with the date of nesting onset (Z = 7.45; p < 0.01), with individuals that commenced nesting earlier in the season producing more nests (Fig. S3), and the number of nests was positively associated with the number of hatchlings produced (Z = 9.16, p < 0.01; Fig. S4). The overall mean nesting interval was 11.4 days, and ranged from 11.1 days in Season 1 to 11.8 days in Season 2, with relatively more observations of 13 days in Season 3 (Fig. S5). Nesting intervals also varied between females, with the mean interval per female ranging from 10 to 16 days after accounting for assumed missed nesting events (>17 days between nests). Based on observations, the longest individual nesting season across the study period was 98 days for a turtle in Season 3 (A18), who laid a total of 9 nests every 12.3 days (S.D. = 1.6 days).

### Emergence success, nest temperatures, and primary sex ratios from empirical nest data

A total of 682 nests were inventoried, with morphometric data taken from a total of 14,057 hatchlings over the course of the four nesting seasons. The overall mean hatchling weight, SCL, SCW, and body depth were 24.3 (S.D. = 2.0) g, 50.0 (S.D. = 2.0) cm, 39.3 (S.D.: = 1.9) cm, and 19.8 (S.D. = 1.0) cm, respectively. All of the hatchling morphological metrics differed statistically among seasons (Fig. S6, Table S2), with all post-hoc pairwise comparisons typically grouping Seasons 1 and 2 together, and 3 and 4 together, except for body depth, for which Season 3 was significantly different from all other Seasons. The mean emergence success across all seasons was relatively high (82.9%), with slight variance across nesting seasons (Fig. 1). Nesting Season 4 had the highest emergence success (84.1%) and Season 2 had the lowest (79.0%). Within seasons, emergence success was not correlated with nesting date, with nests of low emergence success produced sporadically across the nesting season (Fig. 1A). Emergence success was significantly negatively correlated with both maximum and mean temperatures in nests where empirical temperature data were available (Fig. S7), although these regressions only explained a very small proportion of the variation in emergence success (19% and 9%, respectively) and were subsequently not used for downstream analyses. Emergence success varied by individual, with some females producing nests with consistently high success, and others with consistently low success (Fig. S8). Observed nest temperatures varied substantially within seasons, but were relatively similar across nesting seasons (Fig. S9). Nest temperatures for Season 1 were ∼0.5-1.0°C warmer than the three other seasons, however, measurements in that season only had data from the start of the nesting period compared to all other seasons due to the COVID-19 pandemic.

**Figure 1.**
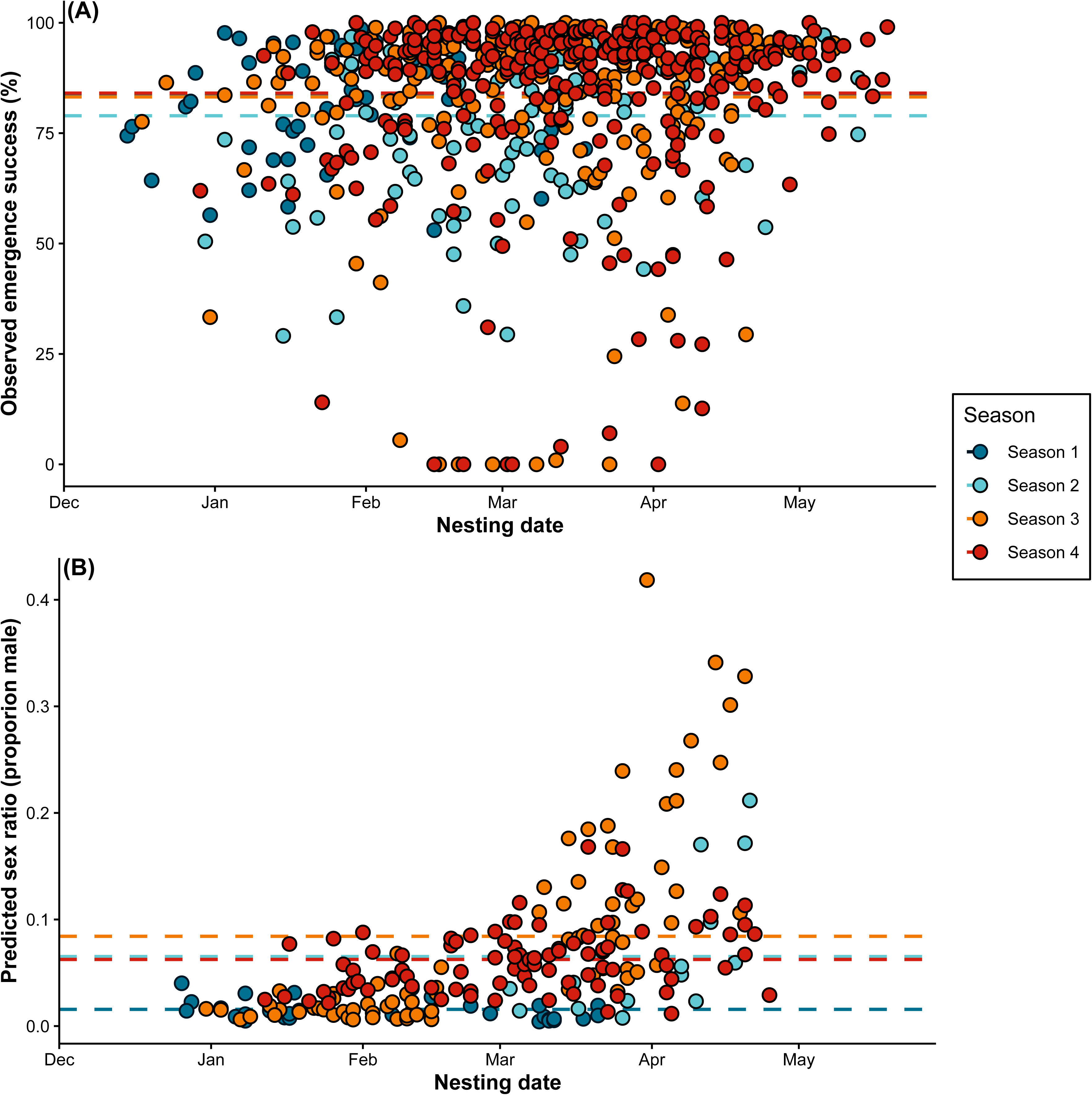
(A) Observed emergence success for 682 green turtle (*Chelonia mydas*) nests distributed across four nesting seasons at Praia do Leão, Fernando de Noronha, Brazil (2019-2023). (B) Modelled primary sex ratios (proportion male) for empirically measured nest temperatures across the four nesting seasons. Dashed lines represent the mean values for the corresponding nesting season. Note that Season 1 was impacted by the global COVID-19 pandemic, and substantially limited sampling.

Predicted sex ratios across all empirically measured nests for the study period were substantially female-skewed, with a mean primary sex ratio of 93.7% female across all seasons. While the predicted sex ratio was consistently female-biased, there was also some variation among nesting seasons, ranging from 98.4% female in Season 1 (noting that sex ratios could not be estimated in the latter half of the season) to 91.6% females in Season 3. Within seasons, sex ratios were relatively consistent across time for nesting Seasons 1 and 4 but showed a substantial increase in male production after March in nesting Seasons 2 and 3 (Fig. 1B).

### Primary sex ratios of repeat nesters

We recorded nest temperatures of between two and seven nests for 39 individual nesting females over two seasons (Seasons 3 and 4). Overall, individual turtles laid nests that produced relatively similar, yet variable sex ratios, with strong variation between individuals (Fig. 2). Sex ratio differed significantly among females in Season 3 (F_19,44_ = 4.42, p < 0.001), but not significantly in Season 4 (F_18,51_ = 1.70, p = 0.07). Similar to the observed trend across all nesters, there was a higher proportion of male offspring produced in the later part of the season by the repeat nesters. In Season 3, females that nested towards the start of the season typically produced nests with less than 10% male offspring, however, this trend shifted for females that commenced nesting after March, with a wider range of sex ratios produced later in the season. Post-hoc tests showed that several females nesting after March had significantly more balanced sex ratios than those at the beginning of the season (Fig. S10). Three females produced nests with a range of 20% between their highest and lowest male production, with A22 producing nests of between 3.6% and 34.1% male hatchlings. Sex ratios were generally more female-skewed in nests laid during Season 4, with very few nests expected to produce more than 10% male offspring. In Season 4, one female (B38) produced nests ranging from 1.2% male to 12.4% males, while B28 produced relatively high male sex ratios (7.8-16.6%) across the three nests laid that year. The only significant pairwise differences in sex ratio in Season 4 was between B28 and three females that nested at the start of the season (B1, B5, and B9).

**Figure 2.**
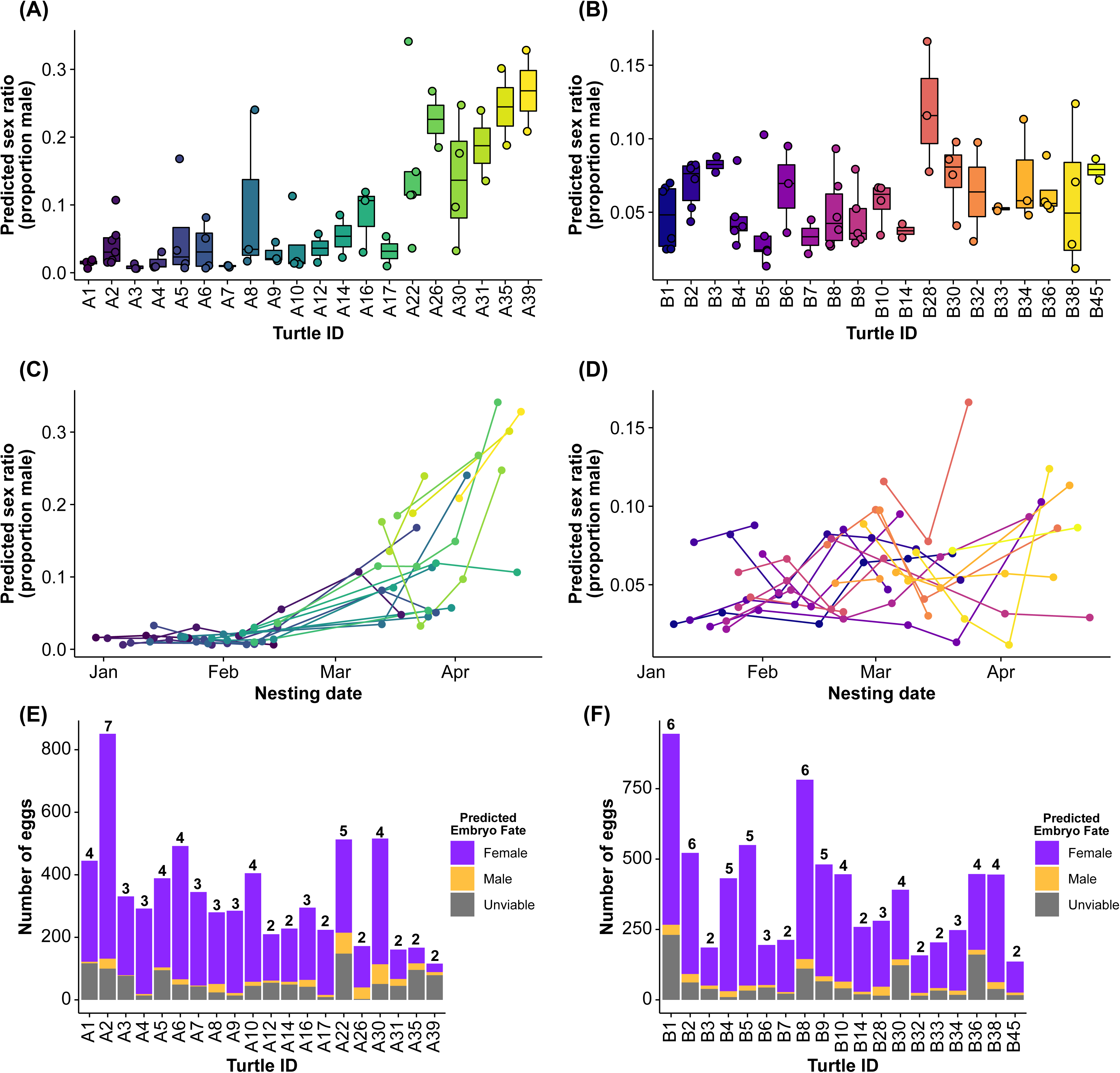
Predicted sex ratios from empirical nest temperatures of repeated nesting females for Seasons 3 and 4. Boxplots of predicted sex ratio for all nests of repeated nesting females for Season 3 (A) and Season 4 (B). (C, D) Predicted sex ratios for each nest plotted against observed nesting date. Colors represent unique individual nesting females, with colors corresponding to turtle ID in (A,B). (E, F) Predicted fate of total aggregate embryos for each female from nests where sex ratio was predicted through embryological models, and emergence success was observed; numbers above bars show total number of nests with temperature recordings per individual female.

When considering all nests laid by our 39 focal nesters, and accounting for observed emergence success, the total number of hatchlings produced varied considerably (Fig 2E, F). In Season 3, individual A2 produced over 800 viable offspring for 7 nests, with the majority of these predicted to be female. Similarly high numbers were produced by B1 and B8 (6 nests each) in Season 4, again with highly female-skewed sex ratios. Although our sex ratio predictions suggest higher male production when nesting occurred later in the season, particularly in Season 3, this pattern did not translate into higher overall male production, because it coincides with fewer nesting events and relatively higher hatchling mortality (Fig. 2E, F).

### Predictions from microclimate models

Metabolic heat raised nest temperatures by up to 5 °C compared to modelled temperatures, with a GAM peaking at approximately 2.5 C (Fig. S11). For outputs from metabolic-heat adjusted microclimate models of observed nests in Seasons 3 and 4, sex ratio predictions were similar to those predicted from the empirical nest data (Fig. 3). These models also showed high variation in sex ratios between females, with less female-skewed nests produced by nesters with later onset of nesting in Season 3 (Fig. 3A), and with relatively stable ratios in Season 4 (Fig. 3B). When combined with empirical observations of total eggs and emergence success, total rookery production featured substantial variation among nesters (Fig. 3C), and the total number of nests laid by each female was a strong determining factor on the total number of male hatchlings she produced over a season (Fig. 4D).

**Figure 3.**
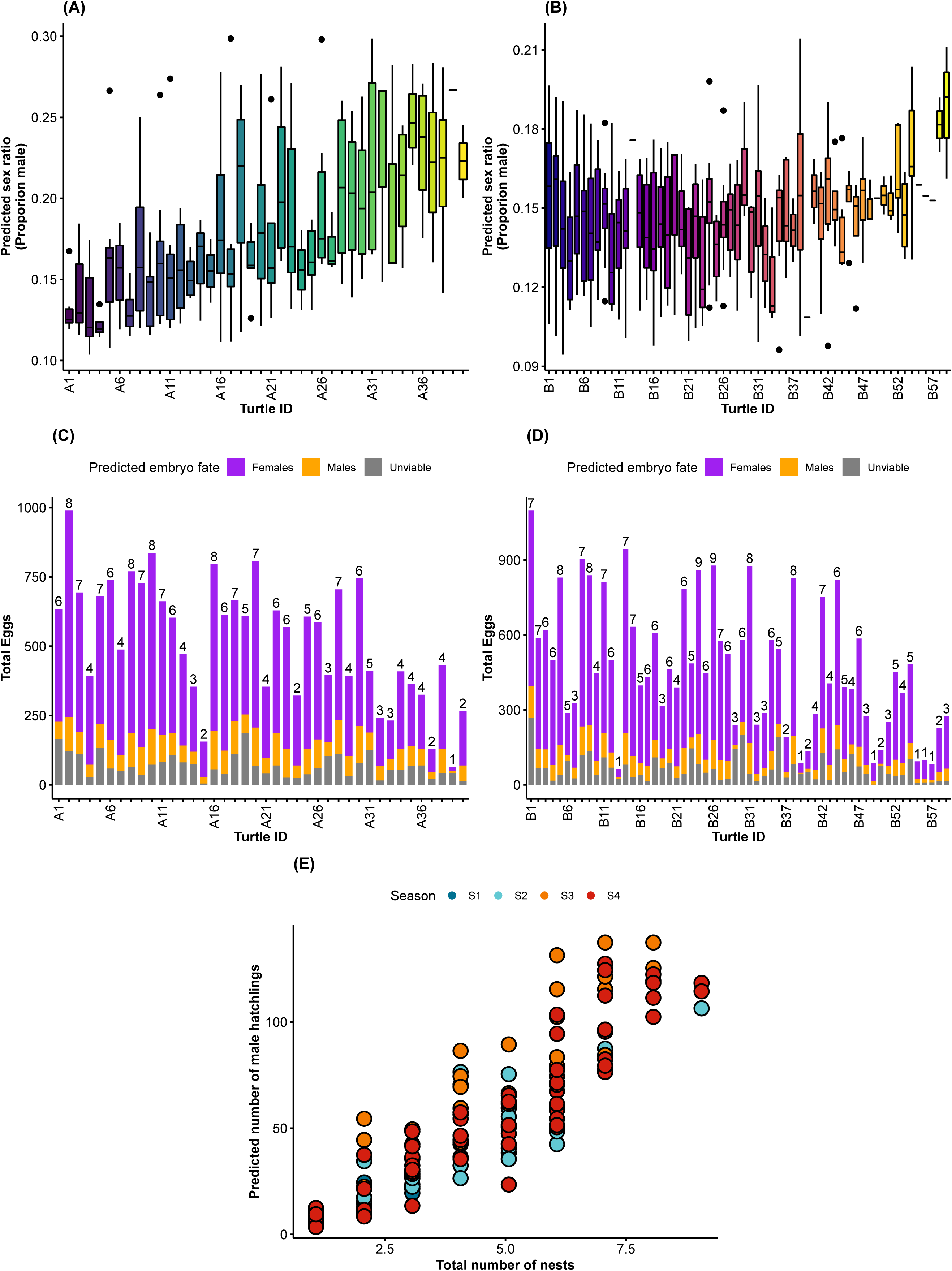
Predicted sex ratios from metabolic-heat adjusted synthetic nest temperature models. (A) Per-female sex ratio across all observed nests for each nester in Season 3 and (B) Season 4. (C) Total predicted per-female hatchling production for Season 3 and (D) Season 4. Numbers above bars represent the total number of nests observed and modelled per female. (E) Regression showing the relationship between total number of nests laid by a nesting female, and number of predicted males contributed to the population across all four seasons (2019-2023).

**Figure 4.**
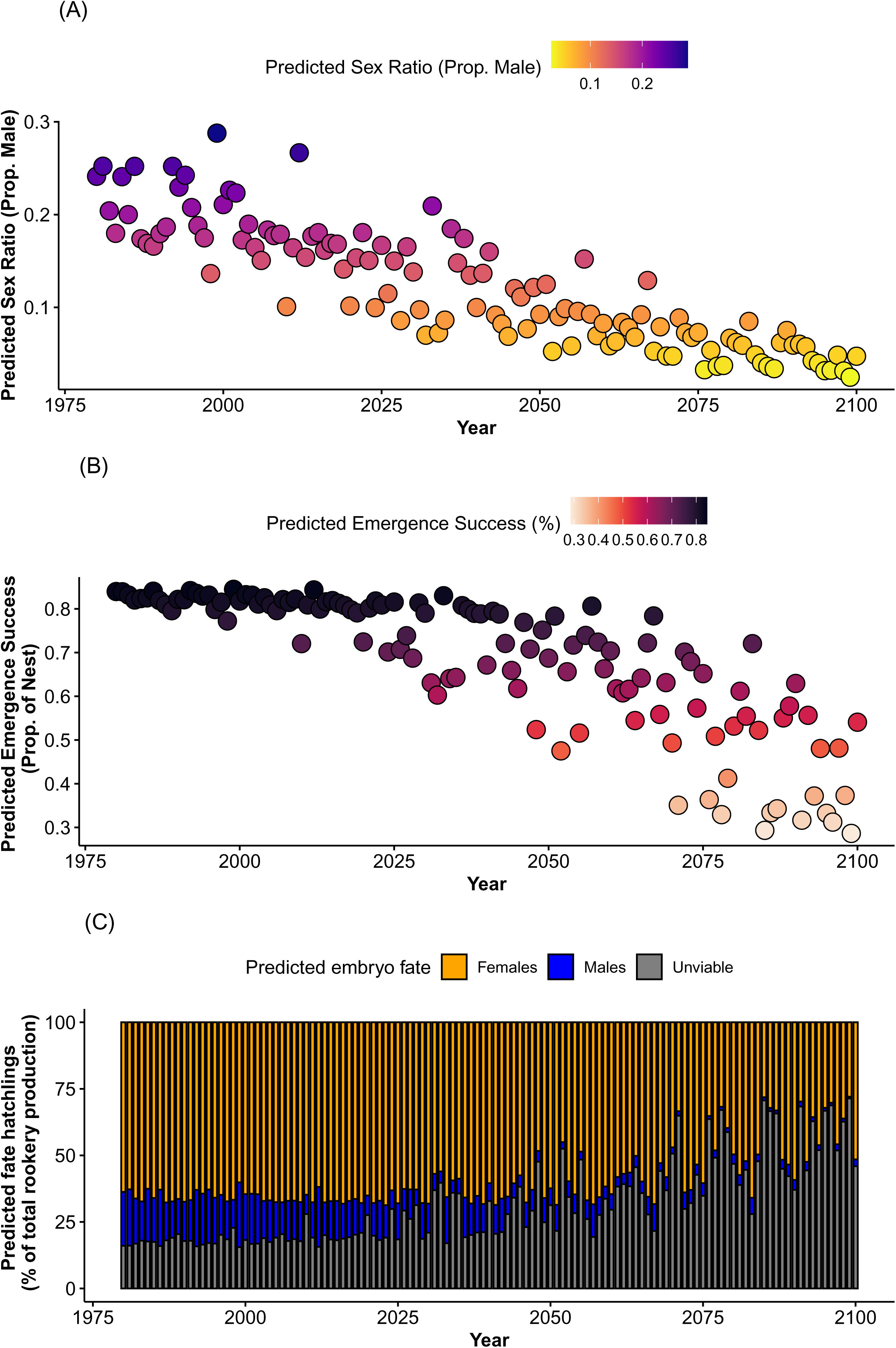
Hind- and forecast emergence success and primary sex ratios from metabolic heat-adjusted microclimate models. (A) Predicted emergence success and (B) sex ratios for all simulated nests from 1980-2100. (C) Combined and nesting distribution-scaled emergence success and sex ratio predictions showing the predicted overall rookery production from 1980-2100.

When accounting for metabolic heat production, overall RMSE and R^2^ values suggested that our models were able to reliably predict nest temperatures at Praia do Leão (Fig. S12). RMSE was typically less than 1° C or less for all seasons. Similarly, R^2^ values were typically greater than 0.80 (Fig S13). Overall, these results suggest that the metabolic-heat adjusted microclimate models were able to provide a relatively high degree of reliability for predicting nest temperatures across our study period.

### Climate change and nesting phenology

Both emergence success and sex ratios are expected to decline substantially over the coming century (Figs. 4A and B) due to increasing nest temperatures. While emergence success has remained relatively constant and high (>80%) since the 1980s, including in recent years, our models suggest a decline to less than 30% by the end of the century, with sharp declines after 2050 (Fig. 4B). Despite our nest temperature models tending to slightly over-predict male hatchlings compared to the empirical observations (see Figs. 2 and 3), our models predicted a strong decline in male production over time (Fig. 4A). Unlike with emergence success, the number of male hatchlings that are being produced has been steadily declining since the 1980s (Fig. 4A), with models suggesting almost no male production by the end of the 21st century (Fig. 4B). When these metrics are combined to generate overall rookery outputs (Fig. 4C), models suggest that in some years towards the end of the century, over 75% of the embryos will perish prior to emergence, with only a small fraction of those viable embryos developing into male hatchlings (Fig. 4C). While there is a degree of sporadic year-to-year variation in these predictions, the overall trend shows a decline in hatchling production at this rookery, and limited male output by the end of the century.

Models of nests laid for every day of the year showed that emergence success and male production are predicted to substantially decrease over the coming century during the current nesting period (Jan-May) (Fig. 5A, B). While there currently exists a period of refuge to maintain a relatively high emergence success (∼May-Oct; Fig. 5B), even at the coolest time of year, nests are expected to produce primarily female offspring at the end of the century (Fig. 5A). Our models suggest that changes in nesting phenology are unlikely to be sufficient to mediate the impacts of climate change on emergence success and sex ratios at this rookery by the end of the century (Figs. 5C, D). In our modelled scenarios, sex ratios are predicted to drop to almost zero in even the most extreme of nesting phenology shifts (Fig. 5C), and emergence success is also predicted to fall below 50%. In order to maintain sex ratios similar to the 2005-2025 average at the end of the century, nesting phenology will need to shift later by ∼100 days (Fig. 5E), and by ∼120 days to maintain contemporary emergence success (Fig. 5F). Interestingly, our models suggest only a small number of years where earlier onset nesting would maintain these levels (n = 6 years for sex ratios and n = 11 years for emergence success), with nesting needing to commence >100 days earlier than current (Figs. 5E, F, S14).

**Figure 5.**
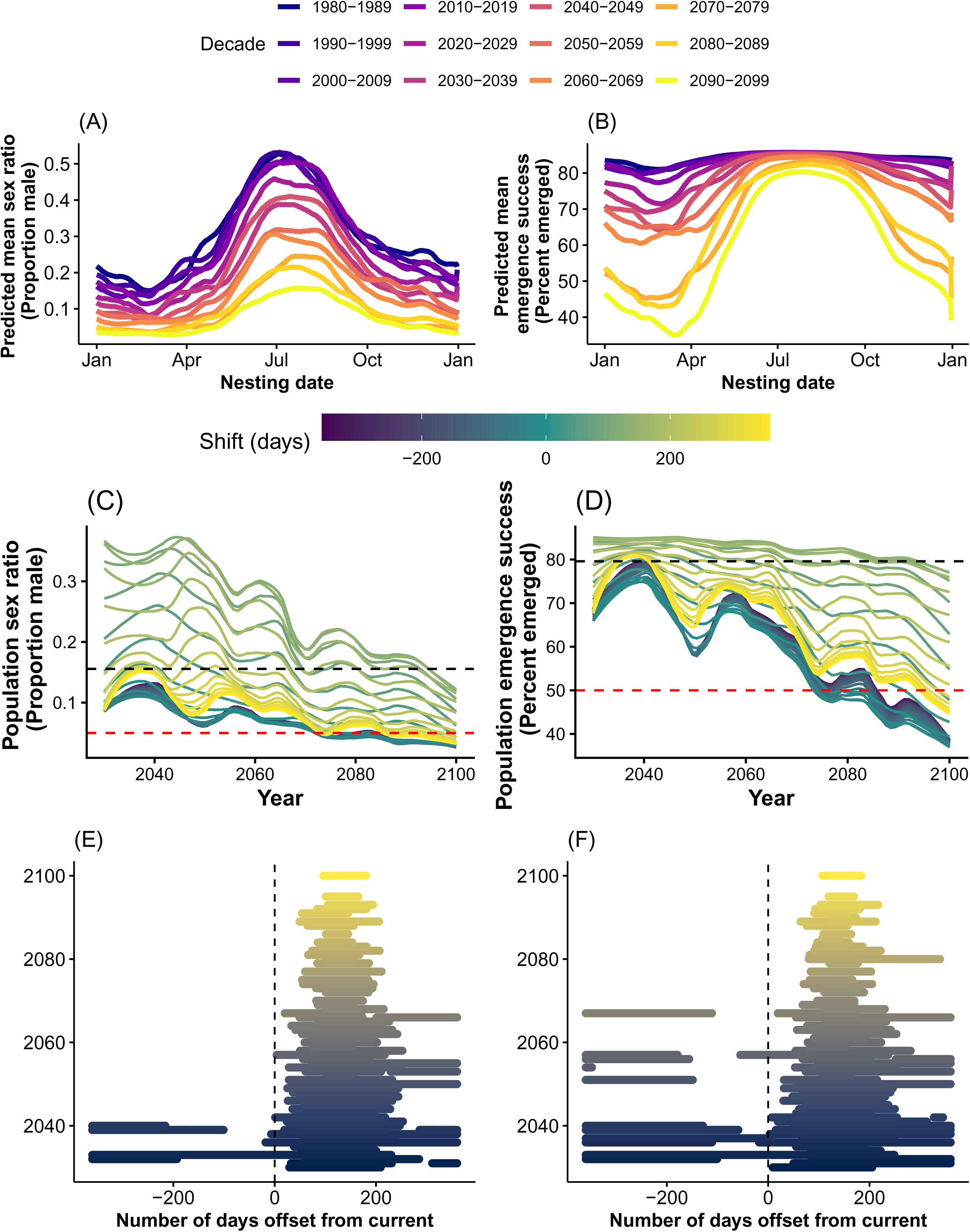
Impacts of nesting phenology shifts on population sex ratios and emergence success. (A) Predicted sex ratio produced for every day of the year averaged across decades from 1980-2100, and (B) predicted emergence success. (C) Predicted population-level sex ratio and (D) emergence success across the nesting season with line color showing the number of days offset (-360 to 360 in 20 day increments, LOESS smoothed regression) from the current nesting distribution. Black dashed lines represent the “current day” mean values (2005-2025), and red dashed lines show 5% male and 50% emergence success. (E) Visual representation of the offset from the current nesting distribution that would result in sex ratios at, or above, mean present day (2005-2025) male production, and (F) mean present day emergence success.

## Discussion

Plasticity in nesting phenology has been proposed as one of the primary mechanisms by which sea turtles may mitigate the impacts of climate warming on embryonic development, yet the extent to which repeated nesting by individual females can buffer hatchling production and primary sex ratios remains unclear. By repeatedly measuring nest temperatures from identified females across entire nesting seasons, we demonstrate that although individual nesters may distribute reproductive effort across seasonal thermal gradients, offspring sex ratios remain relatively consistent within individuals and male production is driven primarily by the onset date of nesting and the total number of clutches produced. Incorporating metabolic heating into mechanistic microclimate models further reveals that phenological shifts alone are unlikely to offset the impacts of projected warming, with male production approaching zero and emergence success declining substantially by the end of the century. Together, these findings suggest that while phenological plasticity provides some resilience to the adverse impacts of climate change, its capacity to buffer reproductive outcomes is limited over the duration of the century.

Through dedicated tracking of individual nesting green turtles at Fernando de Noronha, we were able to directly assess intra-season variation in nest temperature and emergence success, and predict subsequent sex ratios. We show that sex ratios are relatively similar across successive nests of individual turtles, with higher variation between females, especially in Season 3. These results generally reflected population-level trends, with turtles that initiated nesting at a later stage of the season producing nests with a higher male contribution. This is concordant with previous observations in loggerhead turtles (Reneker and Kamel 2016), which demonstrated high variability among females, yet consistent sex ratios within females (however, using incubation duration as a proxy). Similar to previous observations (e.g., Broderick et al. 2003, Lamont and Fujisaki 2014), we show that more nests per individual are correlated with higher fecundity (indicated by overall number of eggs). Interestingly, our models also predict that the total number of males produced is correlated with nest number, with more nests producing more males. Given that the total number of nests produced is negatively correlated with the date of nesting onset, these results suggest that male production is achieved through a combination of many nests over the season or a later onset of nesting. Given that rising temperatures have been linked to changes in nesting phenology in sea turtles (see Patrício et al. 2021), reductions in total nest numbers (Mazaris et al. 2009, Reina et al. 2009, Patel et al. 2016) and nesting intervals (Robinson et al. 2022, Fuentes-Tejada et al. 2025), these two factors are likely to interact under future conditions to shape overall rookery production.

Shifts in nesting phenology are often considered an adaptive strategy for migratory species under changing climates (Patrício et al. 2021). Previous research has shown evidence for nesting phenology shifts in response to rising temperatures in sea turtles (e.g., Weishampel et al. 2004, Pike et al. 2006, Patel et al. 2016, Almpanidou et al. 2018, Witt et al. 2025, Kirkham et al. 2025, Colman et al. 2025), with estimates in the magnitude of shifts ranging between and within species. Despite this assumption, recent models (Laloë and Hays 2023, Fuentes et al. 2024) have predicted that shifts in nesting phenology that align with observed rates of shifts will be insufficient to effectively reduce nest temperatures at global rookeries. Our results align with these models, with even the most extreme shifts in nesting phenology still resulting in warmer nests by the end of the century. Of further concern is that our models show that the most effective shifts in phenology are associated with delays in nesting, rather than earlier onsets. Evidence for climate-driven shifts in nesting phenology in sea turtles has generally documented earlier nesting in response to warming temperatures, particularly in subtropical and temperate populations where seasonal thermal variability is high (Weishampel et al. 2004, Pike et al. 2006, Patel et al. 2016, Almpanidou et al. 2018, Witt et al. 2025, Kirkham et al. 2025, Colman et al. 2025). However, delayed onset of nesting has been reported less frequently, though it has been observed in green and leatherback (*Dermochelys coriacea*) turtles when elevated temperatures at foraging grounds delay individual preparation for reproductive migrations (Dalleau et al. 2012, Robinson et al. 2014, Neeman et al. 2015), which may also result in fewer clutches being produced over the season. Such variability in directions of phenology shifts may be explained by individual- and population-level mechanisms acting simultaneously within a population (Rickwood et al. 2025). Our models of climate change impacts simply explore the response at the population-level, with individual-level changes more difficult to predict.

Across modelled future conditions, phenological shifts in nesting alone were generally insufficient to recover sex ratios to those predicted under contemporary conditions, suggesting that phenological shifts at the population level will not be a sufficient mechanism to alleviate the impacts of rising temperatures at this rookery. Patterns of emergence success mirrored those observed for sex ratios, with substantial declines projected under future climate scenarios and with limited potential for recovery through shifts in nesting timing. Notably, declines in emergence success often preceded or coincided with extreme female-skewing of sex ratios, indicating that climate change threatens overall recruitment back to the rookery through multiple, interacting pathways. As such, population viability may be compromised not only by increasingly female-biased hatchling production, but also by reduced overall hatchling output and elevated embryo mortality, compounding demographic risk and eroding future adaptive potential.

The limited effectiveness of phenological buffering at this rookery likely reflects its proximity to the equator (∼4°S), where seasonal variations and fluctuations in ambient temperature are minimal. Under these tropical conditions, shifts in nesting timing will have little capacity to substantially alter incubation temperatures, especially as projected warming is expected to elevate temperatures throughout the year. This results in a limited capacity for adaptation through the onset timing of nesting, with phenology offering little or no scope for mitigation, and only extreme, unrealistic delays in nesting yielding modest improvements in sex ratio and/or emergence success. These findings are consistent with broader analyses indicating that populations closer to the equator have lower adaptive potential through phenological change than those in more temperate regions (Mazaris et al. 2013, Laloë and Hays 2023, Fuentes et al. 2024). At tropical rookeries such as Fernando de Noronha, reductions in nest temperature are more commonly associated with stochastic rainfall events rather than predictable seasonal cooling. While rainfall can temporarily lower nest temperatures, its timing relative to the thermosensitive period is variable, limiting its predictive reliability as a buffering mechanism (Filonov et al. 2025). Consequently, both behavioral and environmental sources of thermal mitigation are constrained in these systems, further reducing the scope for natural compensation under continued warming.

Together, these results demonstrate that phenological adjustment alone is unlikely to prevent increasingly female-biased sex ratios, declining emergence success, and the associated reductions in effective population size at equatorial rookeries such as Fernando de Noronha. Because male production is elevated at the end of the nesting season and phenological shifts offer limited scope for mitigation, conservation management strategies that implicitly rely on behavioral adaptation are unlikely to be effective in these systems. Instead, interventions that directly modify nest microclimate, such as shading, irrigation, or spatial management of nesting habitat, may be necessary to maintain viable sex ratios and recruitment under future climate scenarios (Fuentes et al. 2012). Such manipulative management interventions require substantial consideration and stakeholder discussions, with potentially maladaptive impacts generated through well-meaning conservation practices. Alternative management practices such as incubation at more favorable conditions such as hatcheries or artificial incubators may provide a less demanding practice, and has already been implemented at several sea turtle rookeries across the globe (Fuentes et al. 2012). More broadly, our findings highlight strong geographic constraints on climate adaptation in species with temperature-dependent sex determination, emphasizing that the capacity for behavioral buffering is unevenly distributed across species’ ranges and is likely to be most limited in tropical, low-seasonality environments.

While the results presented here were generated through robust mechanistic microclimate and embryological models, an inherent shortcoming of all sex ratio models in sea turtles is the inability to validate predictions with empirical sex data, as reliable determination of hatchling sex in the field remains logistically and ethically challenging (Fuentes et al. 2017). Sea turtles lack external sexual dimorphism until maturity (Sifuentes-Romero and Wyneken 2020), which occurs at between ∼7 and 40 years of age, depending on species. As such, there are currently only two reliable methods for determining the sex of hatching sea turtles: either through histological examination of the gonad, which requires euthanasia of many individuals, and is not sustainable or scalable; or by rearing hatchlings to a sufficient size to conduct laparoscopic examination of the gonad, a process which requires specialist personnel and is financially burdensome (Tezak et al. 2020b). While several studies have attempted to elucidate non-invasive biomarkers, these are either only effective in post-hatchling juveniles, or have been inconsistent in their efficacy, although several emerging techniques are providing promising potential (Tezak et al. 2020b, Marín-García et al. 2024, Yen et al. 2026). Ongoing research to develop non-invasive biomarkers for identifying hatchling sex are imperative to allow both predictive model validations and empirical estimates of sex ratios *in situ*, while also permitting more robust investigation into variations in sex determining thresholds. Without such methods, conservation informed by unverified modelling methods may result in maladaptive practices and outcomes.

Additionally, the parameters that define the thermal reaction norm for sex determination, critical thermal maxima of embryos, and embryonic development vary between and within species of sea turtle (Wibbels 2003, Howard et al. 2014, 2015, Laloë et al. 2017, Bentley et al. 2020b). While we employed robust modelling techniques here, we were unable to fully parameterize the embryological models with population-specific thresholds due to the absence of data. As such, these models include many assumptions that may not hold true to the absolute outputs of the rookery. Despite these shortcomings, we note that our results show relative changes in sex ratio and mortality over time, with observed trends likely to hold true even if empirical values vary from predictions.

Together, our results suggest that sea turtles at tropical rookeries, such as Fernando de Noronha, may not possess the ability to effectively alleviate climate change impacts through shifts in nesting phenology. We show that individual green turtles at Fernando de Noronha are able to influence the thermal environment and subsequent sex ratio of their nests through variation in the date that they initiate nesting, with later nests likely contributing relatively more male recruits to this population. However, our models also suggest that even if delayed nesting is possible in this population, the resulting benefits are likely to be lost under future conditions, as year-round warming eliminates windows of high rookery production, potentially threatening the continued persistence of nesting in this region.

## Data accessibility

All data associated with analyses are available on Dryad (DOI: 10.5061/dryad.5dv41nsn8). Scripts associated with all analyses can be accessed through GitHub: https://github.com/bpbentley/FeN_Cmydas_sex_ratios

## Acknowledgements

The authors acknowledge the field support provided by Mariana Ricciardi, Nathalia Rodrigues, Daniele Macedo, Luna Moreno, and Adriana Jardim, as well as additional support provided by Projeto TAMAR, ICMBIO and Centro TAMAR, and Projeto Golfinho Rotador. We also thank Jonathan Monsinjon for assistance with development of the EmbryoGrowth models. Funding for this project was provided through the National Science Foundation (NSF: OCE-1904818, IOS-1904439, IOS-1904615), and all research was conducted under a Florida State University IACUC permit (#PROTO202000076). Models were run using the UMass UNITY supercomputer supported by the Massachusetts Green High Performance Computing Center (MGHPCC).

