## Supplementary Tables and Figures for "Phenological plasticity provides limited resilience to climate warming in a sea turtle species with temperature-dependent sex determination"

**Table S1.** Total number of loggers deployed per season, number of unique nesting females covered, and temporal coverage of nesting period.

| **Season** | **Total number of loggers deployed within nests** | **Number of reliable datasets obtained for analysis** | **Total unique nesting females covered** | **Temporal coverage of temperature loggers** |
| --- | --- | --- | --- | --- |
| 1 (2019/2020)* | 41 | 38 | 24 | 2019-12-20 to 2020-05-28 |
| 2 (2020/2021) | 38 | 16 | 18 | 2021-01-15 to 2021-06-13 |
| 3 (2021/2022) | 166 | 110 | 40 | 2021-12-23 to 2022-07-07 |
| 4 (2022/2023) | 148 | 121 | 52 | 2023-01-11 to 2023-07-11 |

* Deployment of temperature loggers in the 2019/2020 nesting season was interrupted by the global COVID-19 pandemic.

**Table S2.** Hatchling morphometrics across seasons. Differences were tested using generalized mixed models with nest ID as a random effect (trait ~ Season + (1 | nest_ID)).

| **Season** | **Total number of hatchlings measured** | **Mean weight (g)** | **Mean straight carapace length (SCL; mm)** | **Mean straight carapace width (SCW; mm)** | **Mean body depth (BD; mm)** |
| --- | --- | --- | --- | --- | --- |
| Season 1: 2019-2020 | 995 | 23.23 (1.99)^A^ | 49.13 (1.72)^A^ | 38.72 (1.74)^A^ | 19.30 (1.00)^A^ |
| Season 2: 2020-2021 | 3,561 | 23.93 (1.81)^A^ | 49.26 (1.80)^A^ | 38.48 (1.73)^A^ | 19.55 (0.91)^B^ |
| Season 3: 2021-2022 | 5,257 | 24.43 (1.99)^B^ | 50.49 (1.94)^B^ | 39.88 (1.96)^B^ | 20.11 (1.07)^A^ |
| Season 4: 2022-2023 | 4,244 | 24.68 (1.89)^B^ | 50.31 (1.95)^B^ | 39.53 (1.71)^B^ | 19.67 (0.96)^A^ |
| GLMM ANOVA F-value |  | 6.62 | 19.11 | 22.97 | 16.18 |
| GLMM ANOVA p-value |  | <0.001 | <0.001 | <0.001 | <0.001 |

N.B. letter superscripts represent pairwise statistical significance, as tested through Tukey post hoc tests, where differing letters represent significance at 0.95 alpha level. Letters are distinct between measured traits.

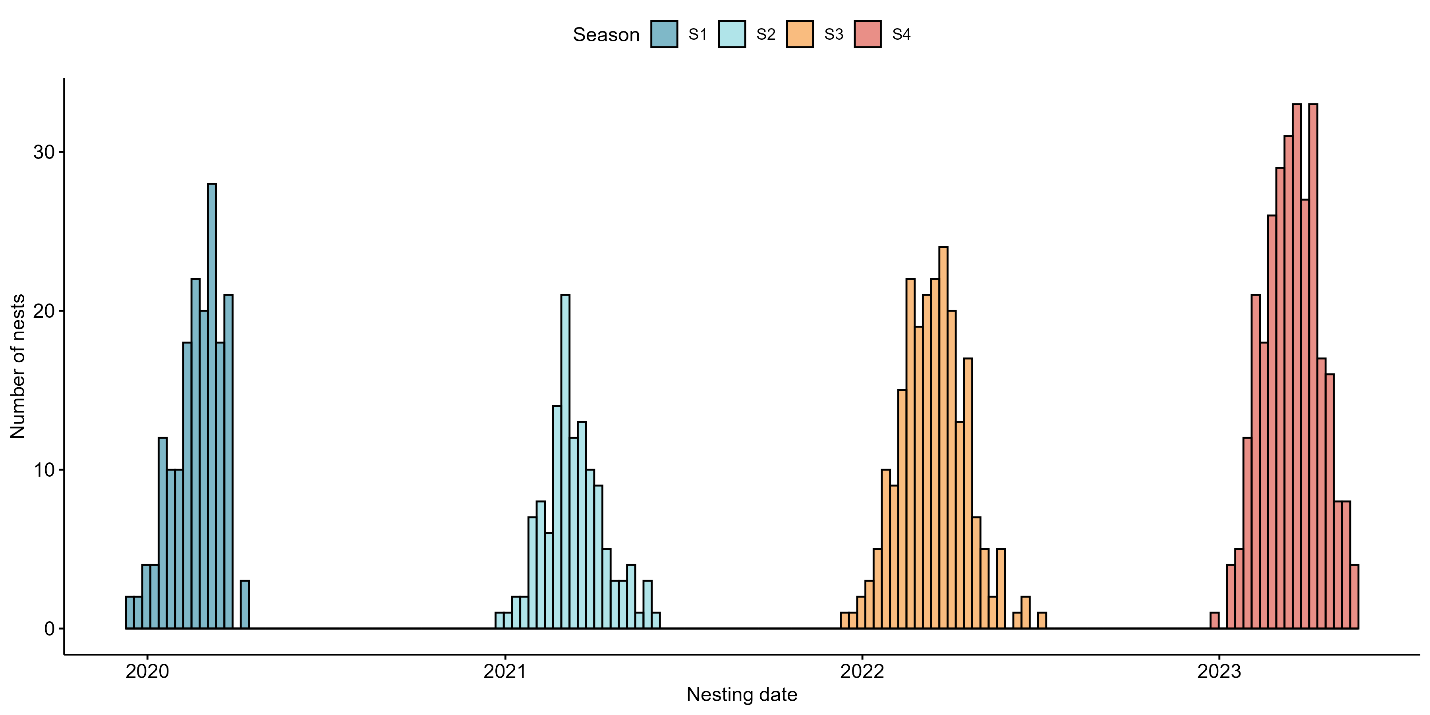

Fig S1. Nesting distribution of green turtles across four nesting seasons (2019-2023) at Fernando de Noronha, Brazil. Nesting typically commenced in mid- to late-December, and concluded in late May/early June. Note that observations for Season 1 are truncated due to the COVID-19 pandemic.

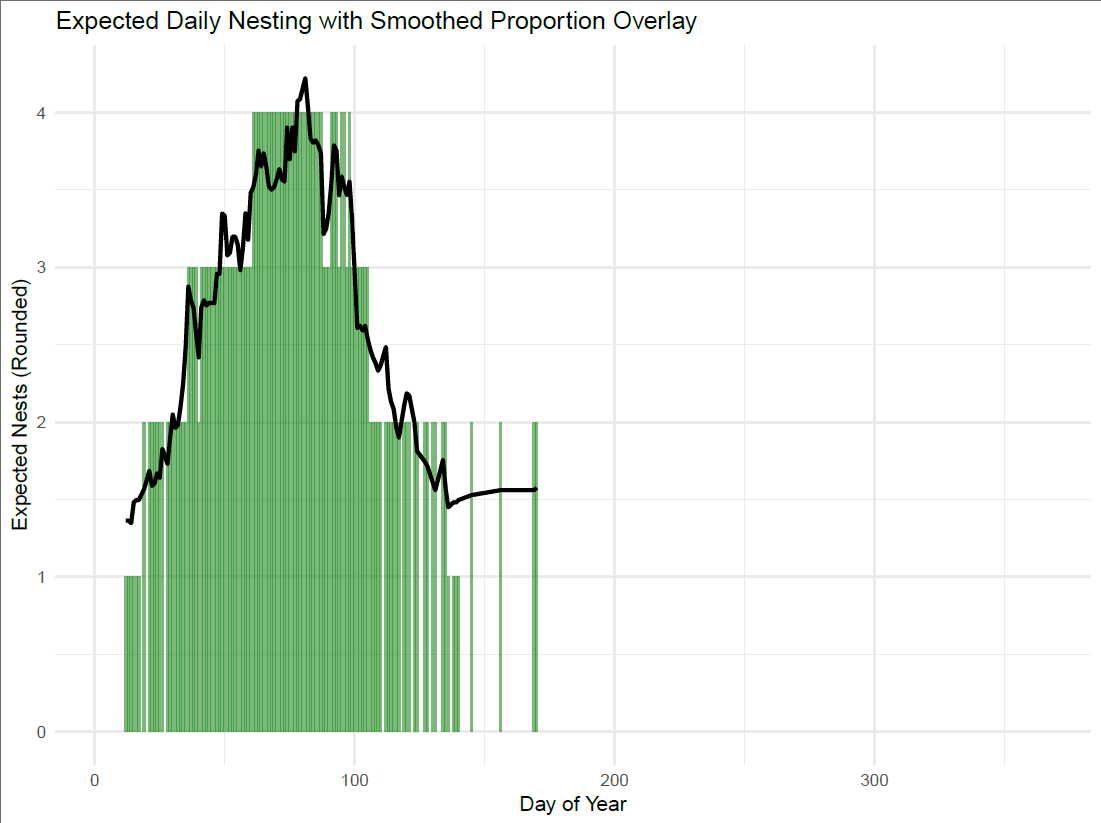

**Figure S2**. Smoothed, 10-day rolling average of nest number across two full nesting seasons (Seasons 3 and 4; black line) and corresponding number of nests per day of year used in all sex ratio and emergence success predictions (green bars; total = 334 nests).

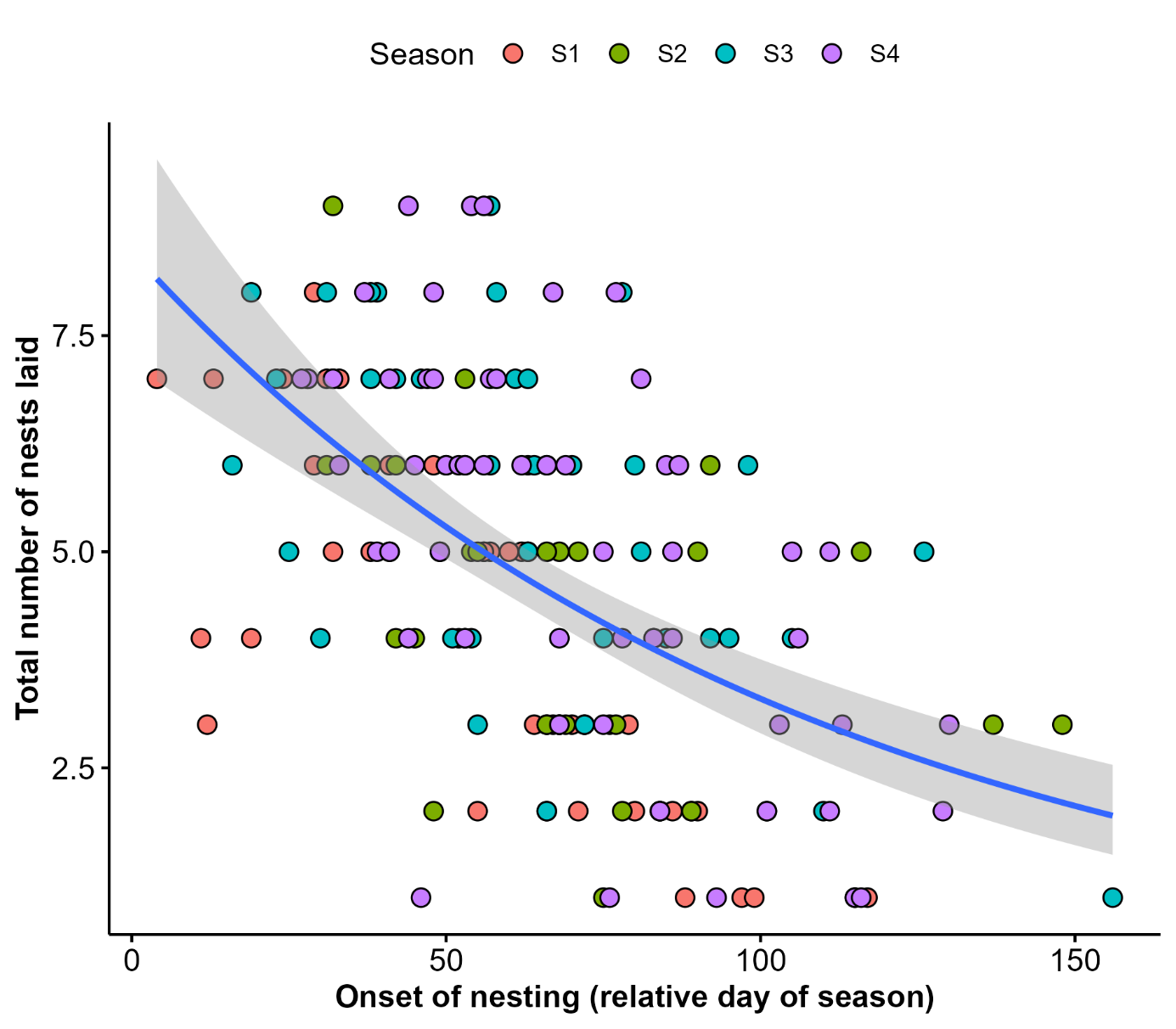

Fig S3. Relative date of nesting onset has a negative relationship with the total number of nests laid for each individual nesting turtle.

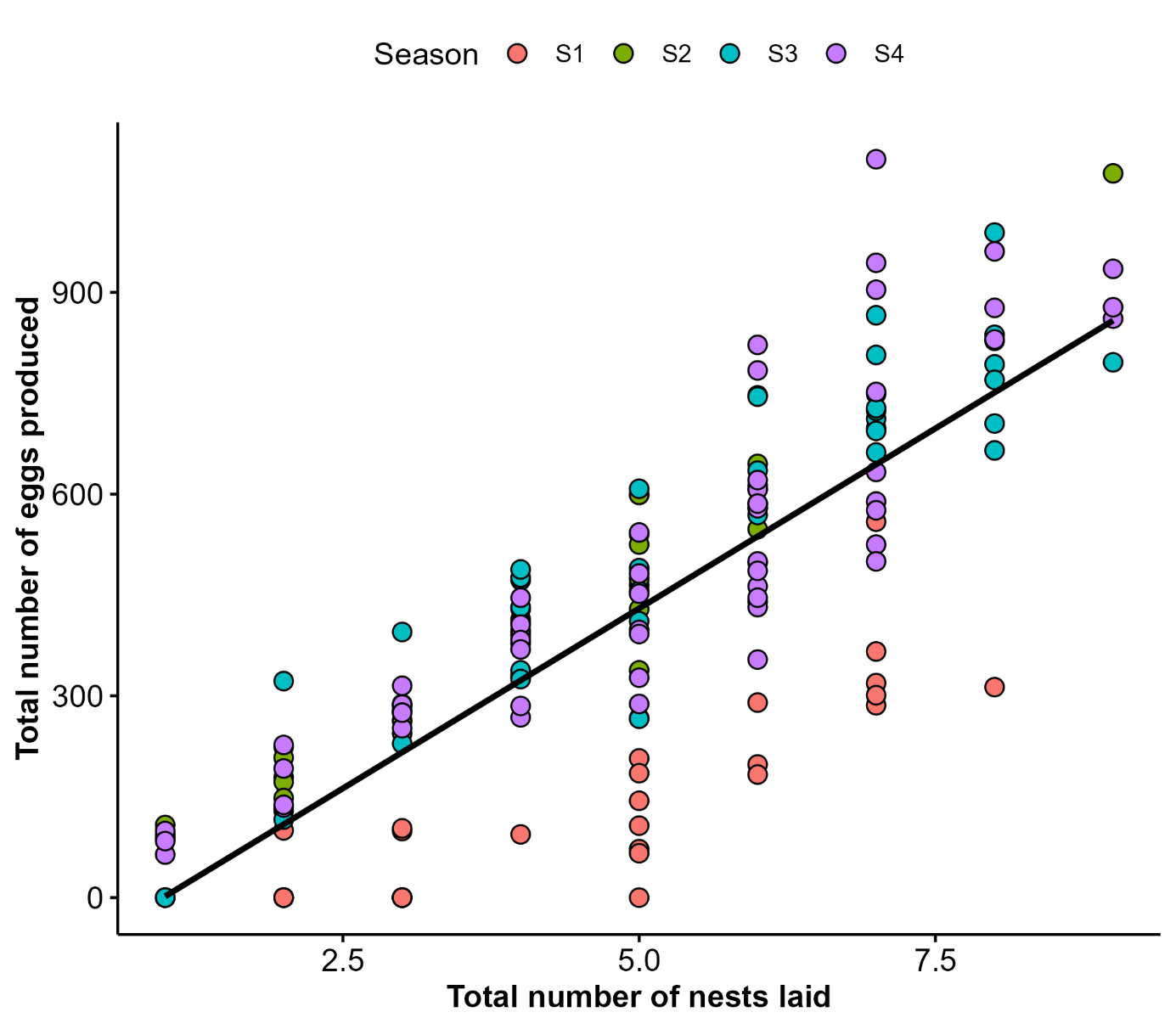

Fig S4. Total number of nests laid by an individual turtle was positively correlated with total number of eggs produced. Note that Season 1 was truncated by the COVID-19 pandemic.

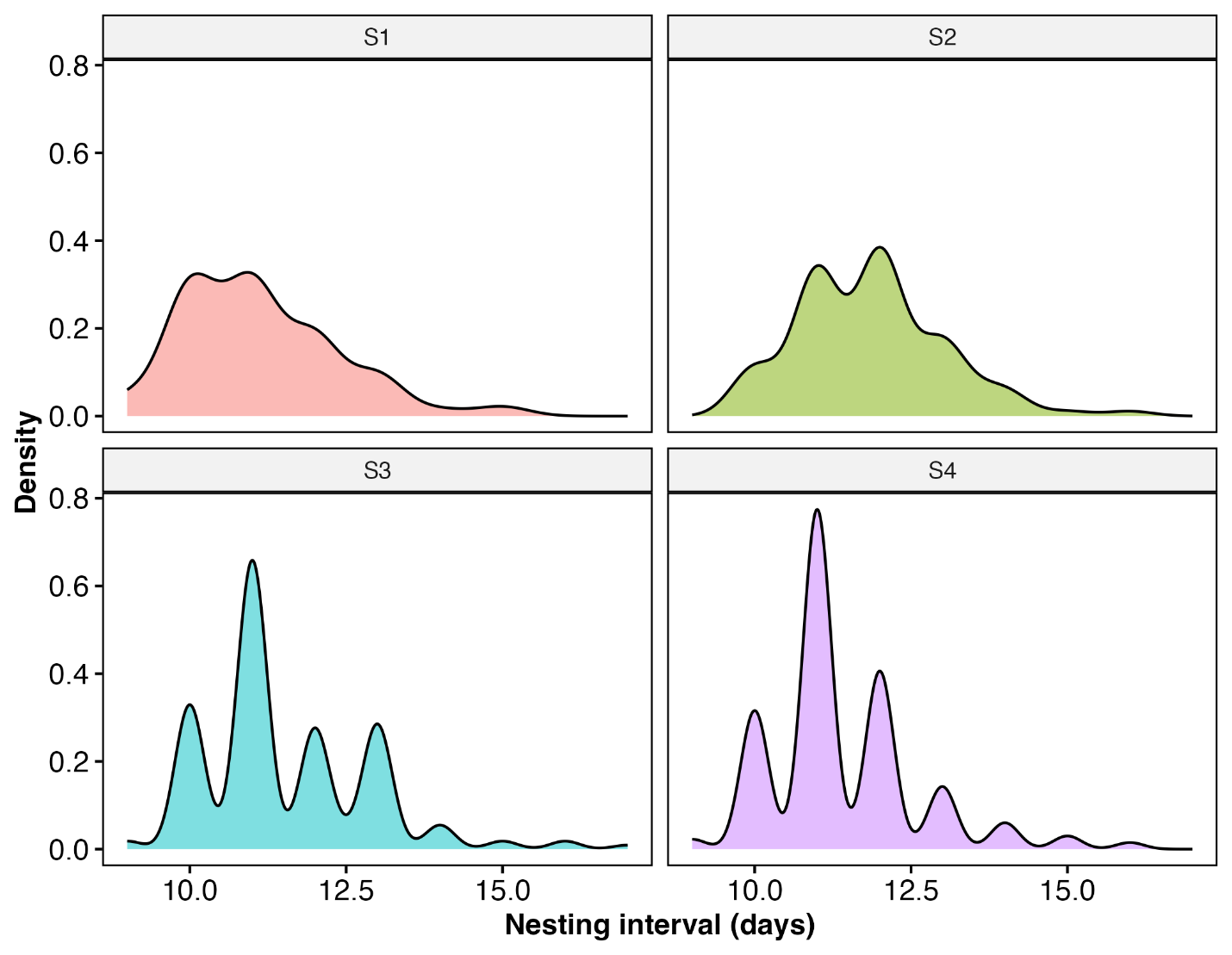

Figure S5. Density plots showing the frequency of nesting intervals (9-16 days) between seasons.

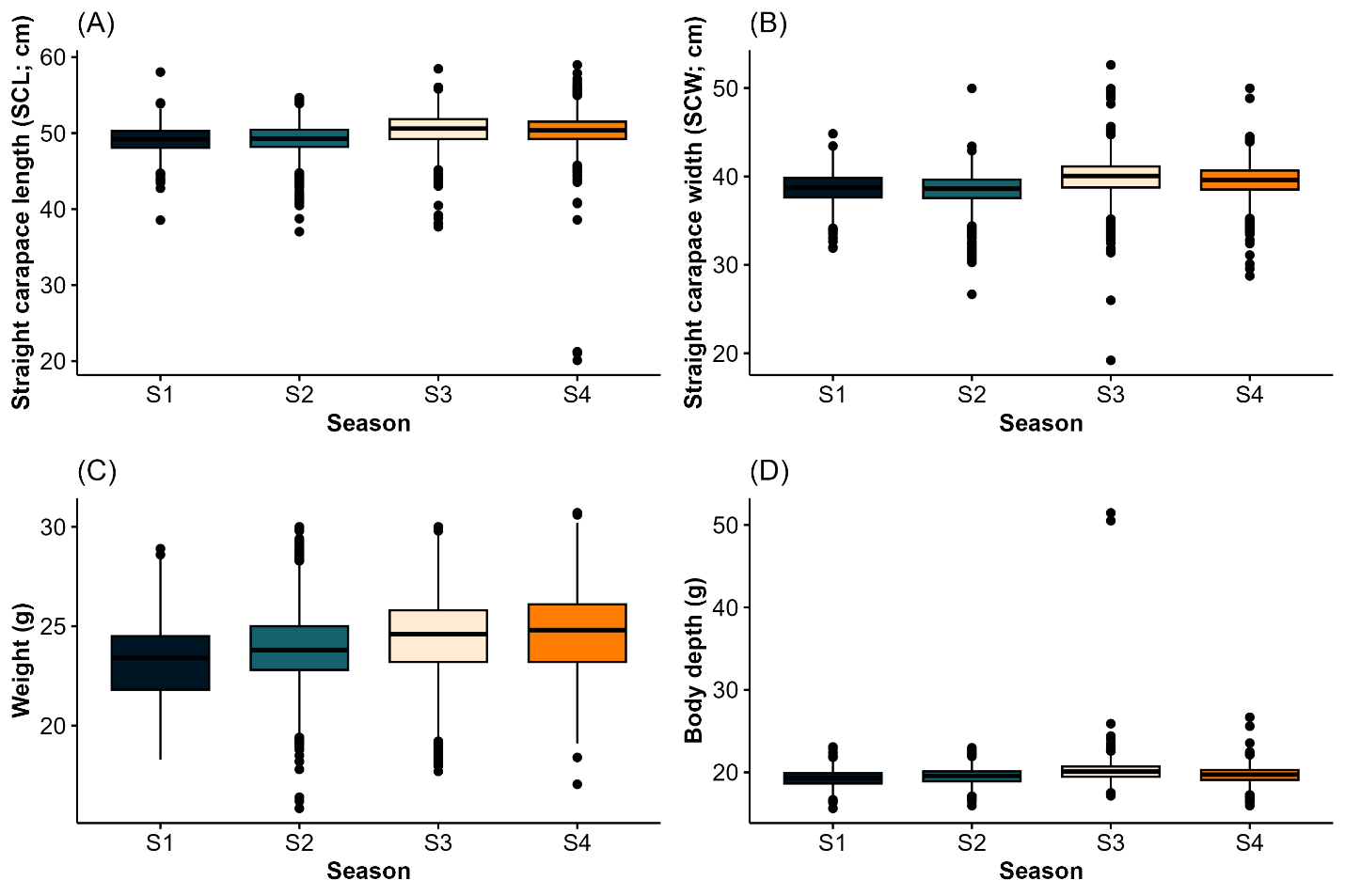

**Figure S6.** Boxplots showing season-wide morphometrics. Panels represent (A) straight carapace length (SCL); (B) straight carapace width (SCW); (C) weight; and (D) body depth. Outliers (+/- 2 S.D. from mean) were removed prior to analyses.

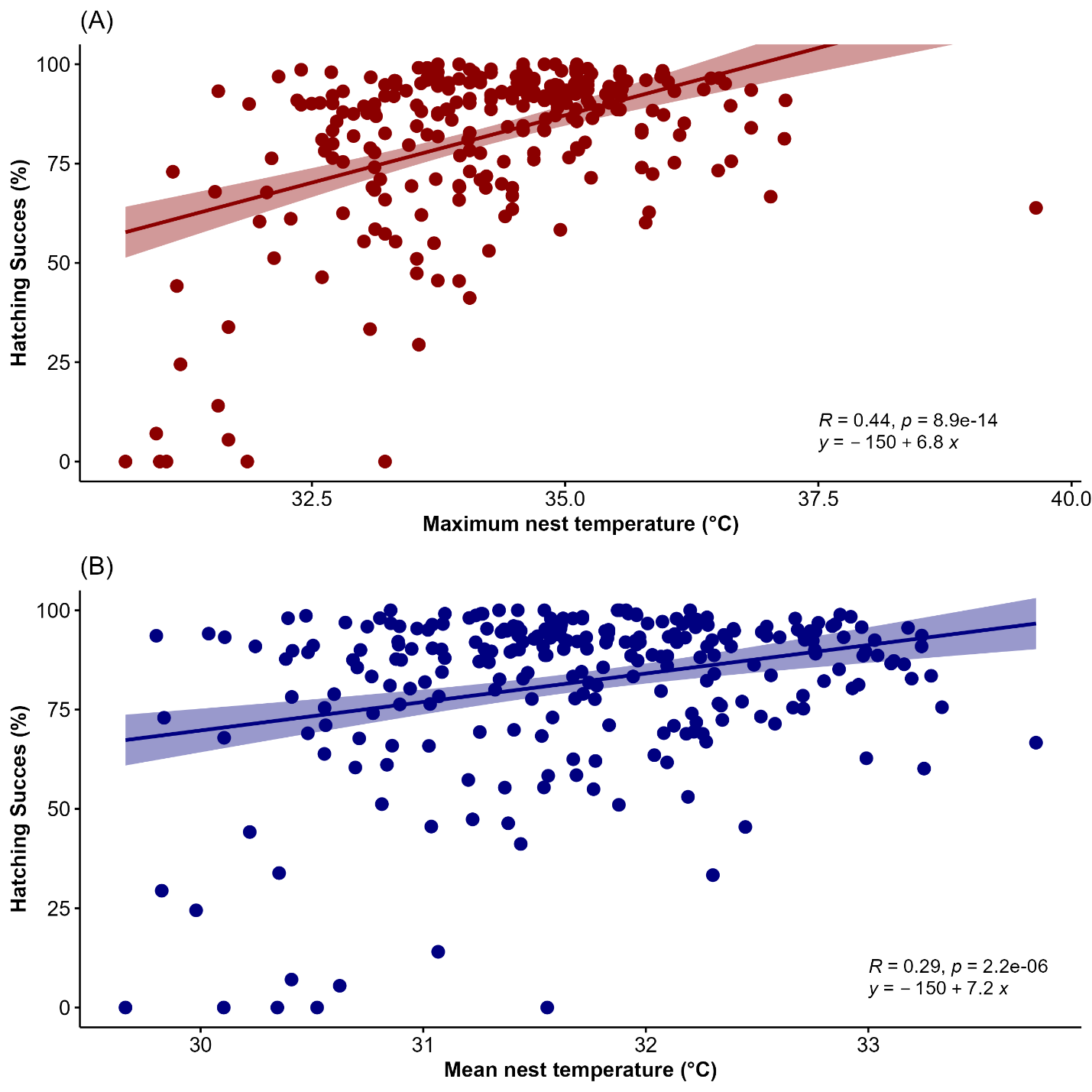

**Fig S7.** Relationship between hatching success and (A) maximum nest temperature; and (B) mean nest temperature through incubation. Trend lines represent linear regressions, with 95% confidence intervals shaded.

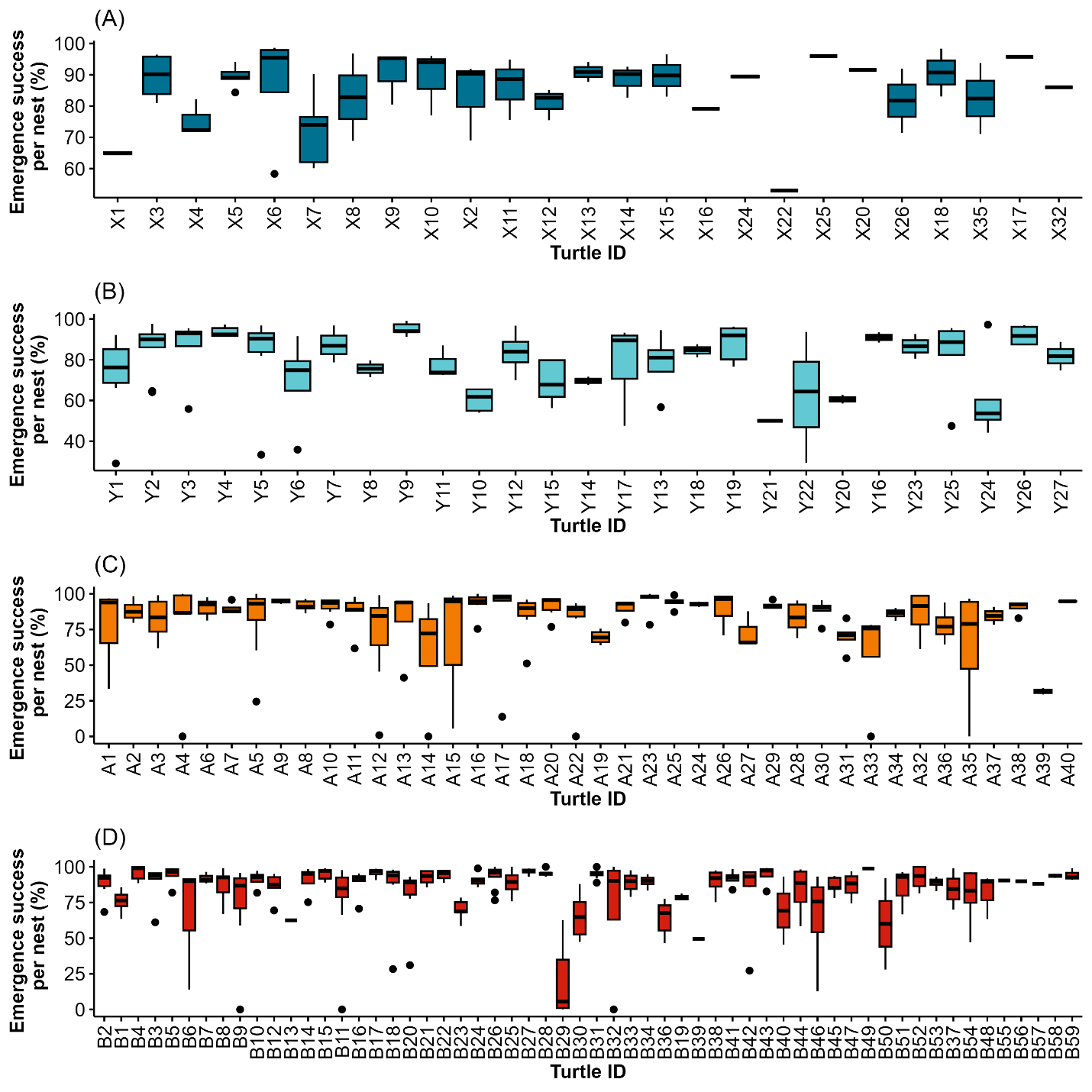

Figure S8. Female-specific emergence success for individually identified turtles across the four seasons at Fernando de Noronha (2019-2023). (A) Season 1, (B) Season 2, (C) Season 3, (D) Season 4.

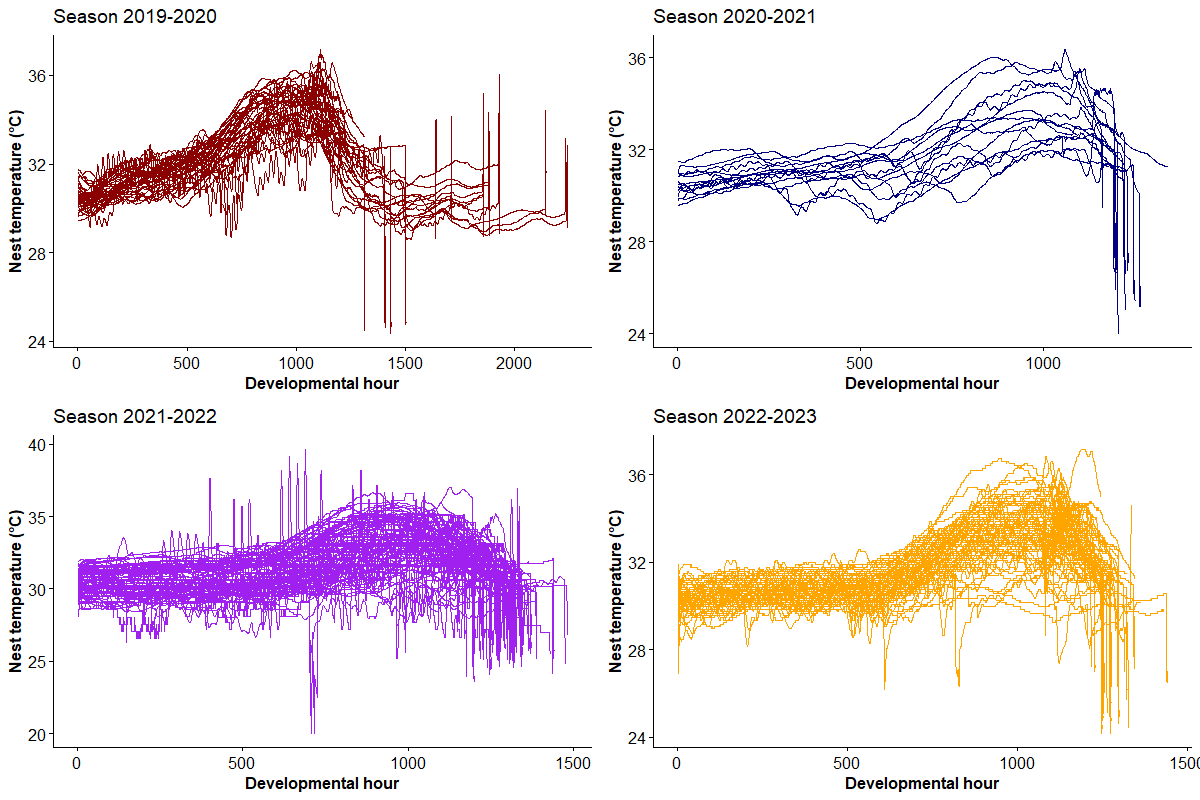

**Figure S9.** Observed nest temperatures across four seasons at Fernando de Noronha, scaled per hour of development.

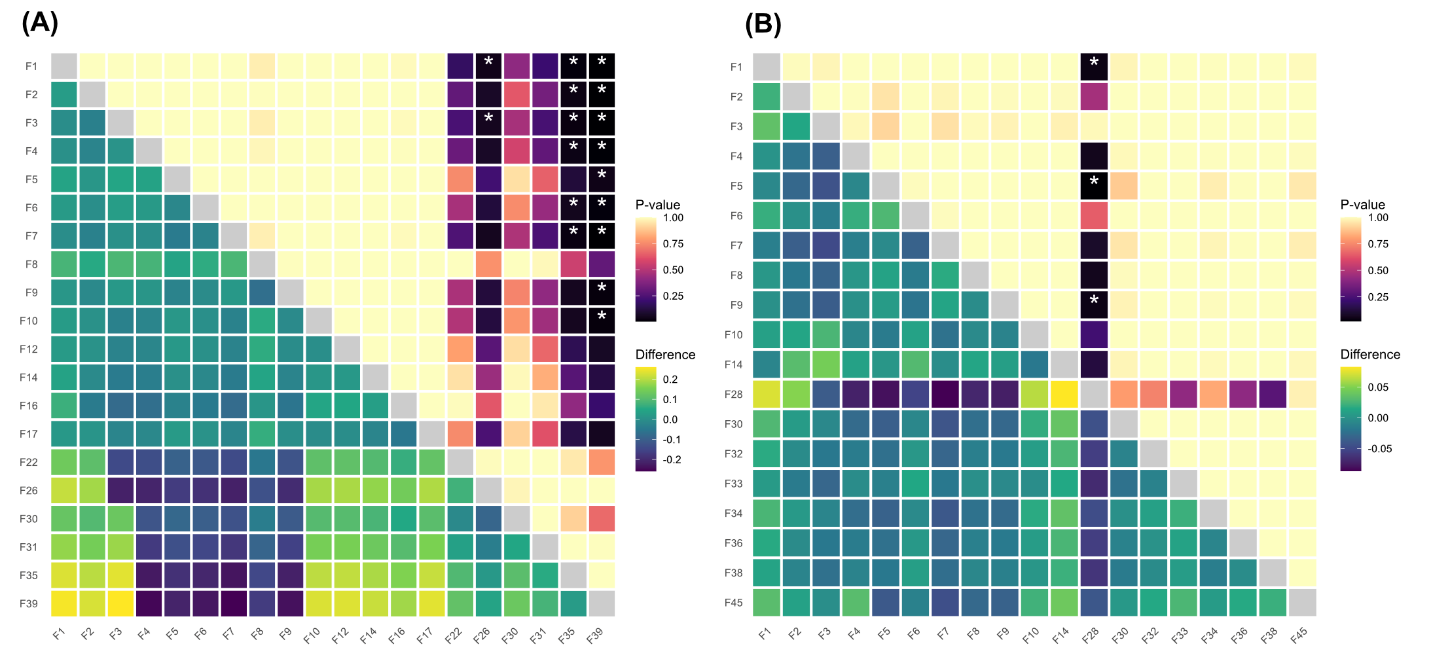

**Figure S10.** Pairwise comparisons (Tukey post-hoc tests) of predicted sex ratios from empirically measured nests between tracked females across (A) nesting Season 3 [2021/2022] and (B) Season 4 [2022/2023]. Colors on the upper diagonal represent pairwise p-values, while the lower diagonal shows absolute differences in sex ratios. Asterisks represent significant pairwise differences (p < 0.05).

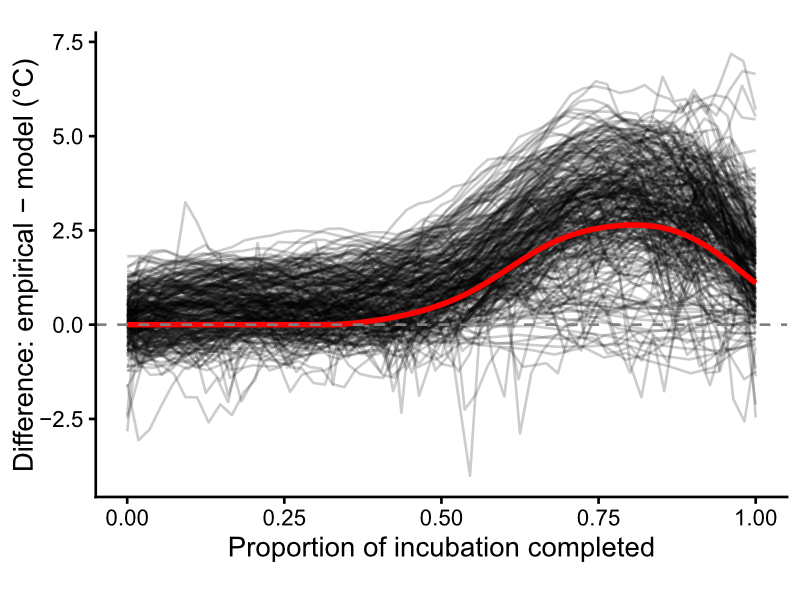

**Figure S11**. Difference between NicheMapR sand temperature model and empirical nest temperatures at Fernando de Noronha between 2019 and 2023 across incubation duration. Each black line represents an individual nest, and the red line shows the model bias-corrected generalized additive model (GAM) which was subsequently added to all sand models for sex ratio and emergence success predictions.

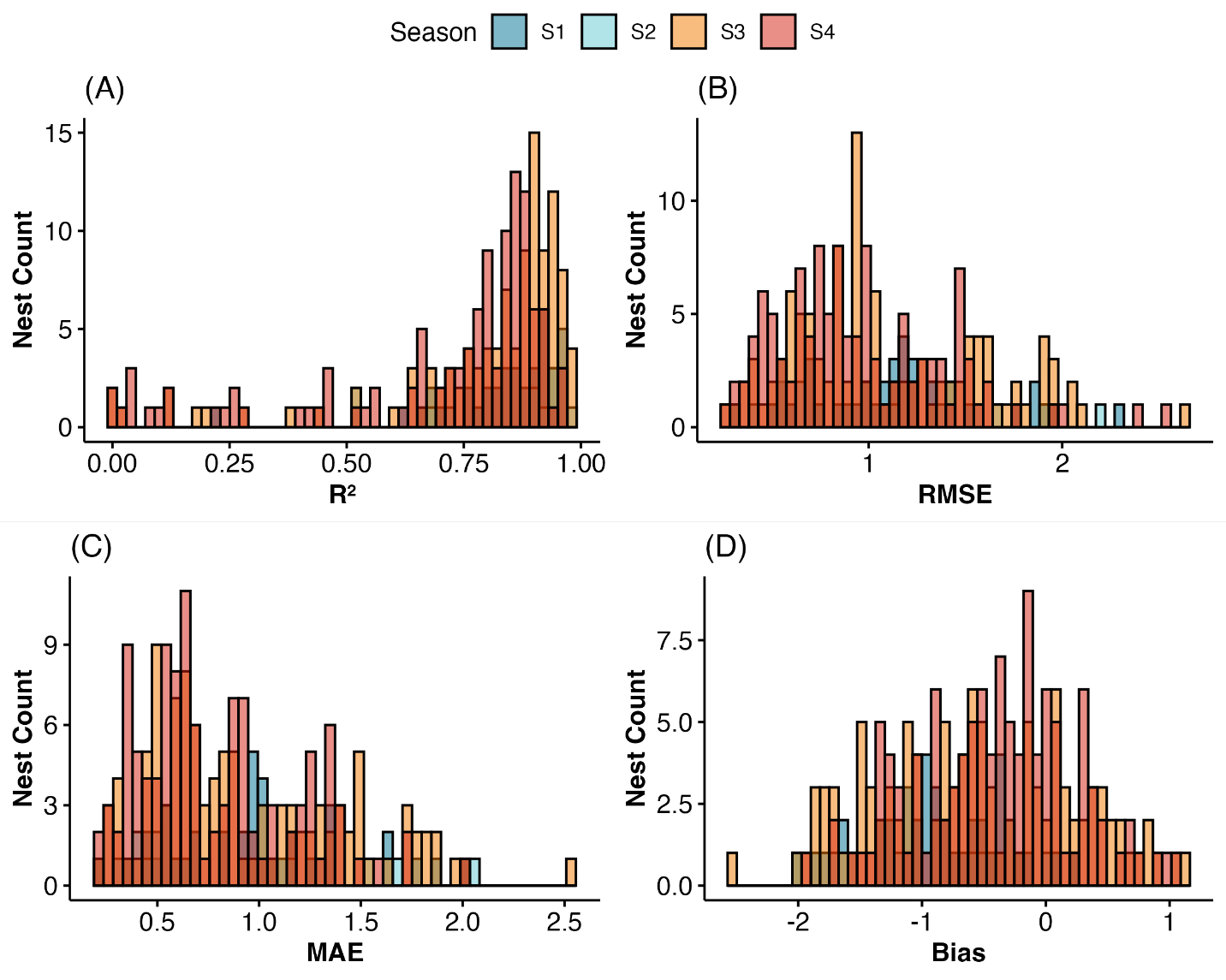

**Figure S12**. Histograms of error metrics for both the metabolic heat-adjusted models across all four seasons. (A) Shows the correlation between observed and modeled daily temperatures; (B) the root-mean standard error (RMSE); (C) mean absolute error; and (D) overall model bias, for each nest. Colors represent models from each season.

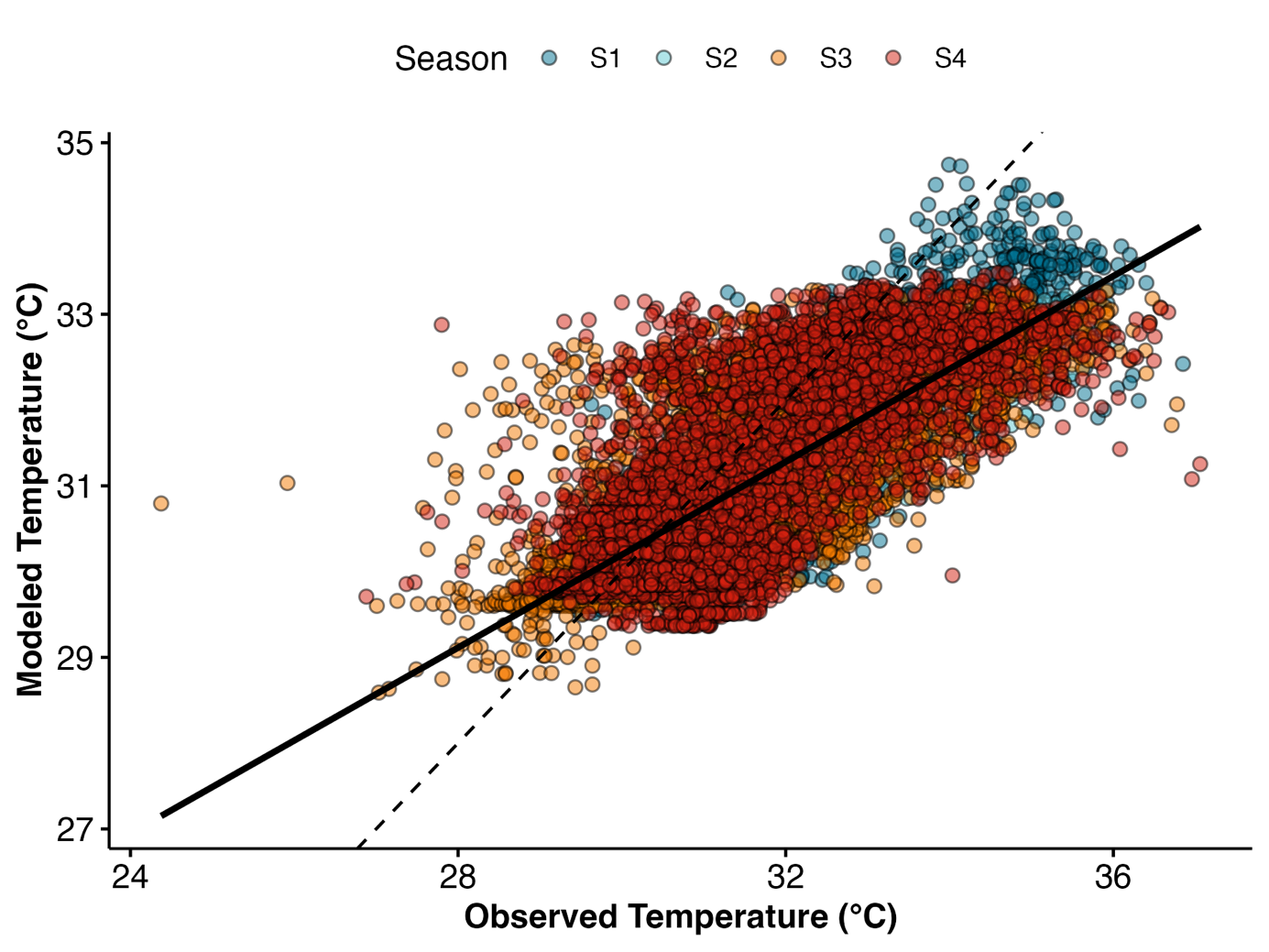

**Figure S13**. Regression between observed nest temperatures and metabolic-heat adjusted modeled temperatures. Solid line shows the regression line, and the dashed line shows a 1:1 regression with an intercept of 0. Colors represent Season.

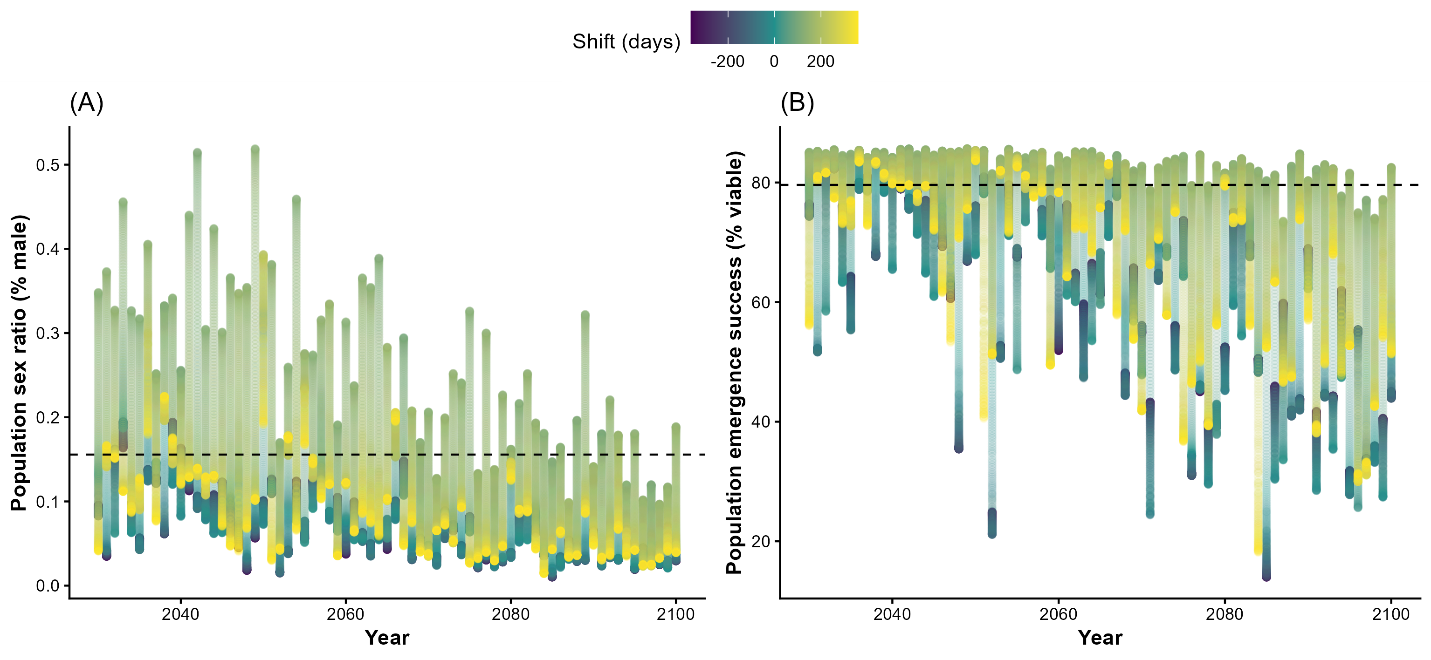

**Figure S14.** Influence of nesting phenology shifts on (A) sex ratios and (B) emergence success from 2030-2100 at Fernando de Noronha. Black dashed line shows the “contemporary” mean values (2005-2025).
